# The *Hypolimnas misippus* genome supports a common origin of the W chromosome in Lepidoptera

**DOI:** 10.1101/2023.03.24.533969

**Authors:** Anna Orteu, Shane A. McCarthy, Emily A. Hornett, Matthew R. Gemmell, Louise A. Reynolds, Ian A. Warren, Ian J. Gordon, Gregory D. D. Hurst, Richard Durbin, Simon H. Martin, Chris D. Jiggins

## Abstract

Moths and butterflies (Lepidoptera) have a heterogametic sex chromosome system with females carrying ZW chromosomes and males ZZ. The lack of W chromosomes in early diverging lepidopteran lineages has led to the suggestion of an ancestral Z0 system in this clade and a B chromosome origin of the W. This contrasts with the canonical model of W chromosome evolution in which the W would have originated from the same homologous autosomal pair as the Z chromosome. Despite the distinct models proposed, the rapid evolution of the W chromosome has hindered the elucidation of its origin. Here, we present high-quality, chromosome-level genome assemblies of two *Hypolimnas* species (*Hypolimnas missipus* and *Hypolimnas bolina)* and use the *H. misippus* assembly to explore the evolution of W chromosomes in butterflies and moths. We show that in *H. misippus* the W chromosome has higher similarity to the Z chromosome than any other chromosome, which could suggest a possible origin from the same homologous autosome pair as the Z chromosome. However, using genome assemblies of closely related species (ditrysian lineages) containing assembled W chromosomes, we present contrasting evidence suggesting that the W chromosome might have evolved from a B chromosome instead. Crucially, by using a synteny analysis to infer homology, we show that W chromosomes are likely to share a common evolutionary origin in Lepidoptera. This study highlights the difficulty of studying the evolution of W chromosomes and contributes to better understanding its evolutionary origins.

**Significance:** Butterflies and moths have a sex determination system in which females carry two different sex chromosomes, Z and W, while males carry two copies of the Z. The evolutionary origin of the W chromosome has been elusive, with many possible scenarios being suggested, such as the independent evolution of W chromosomes in many butterfly and moth species. Here, we present genome assemblies of two *Hypolimnas* butterfly species and use one of them to shed light on the evolution of the W chromosome. We show that W chromosomes across butterflies and moths are very similar which suggests a shared common origin.

## Introduction

Sex chromosomes are highly variable in eukaryotes and have evolved independently multiple times (Bachtrog et al. 2014, 2011; Beukeboom & Perrin 2014). In animals there are multiple types of chromosomal sex determination systems but two are predominant: male heterogamety as seen in mammals where males are XY and females XX; and female heterogamety as seen in birds where females are ZW and males ZZ (Beukeboom & Perrin 2014). These heteromorphic sex chromosomes (XY and ZW) can potentially originate from different processes. One possibility is that they initially arise from a pair of autosomes that evolve genetic sex determination and, through a process of reduced recombination, sex specific mutations accumulate (Wright et al. 2016). This reduction in recombination can also lead to gene depletion and the accumulation of repeat and transposable elements (TEs), which are common characteristics of sex specific (Y or W) chromosomes (Bachtrog et al. 2011; Wright et al. 2016). However, discerning between cause and consequence can be difficult, as repeats and TEs could be enhancing the reduction in recombination rather than resulting from it. Additionally, other autosomes may fuse to the sex chromosomes and become differentiated. Alternatively, the W/Y chromosomes can originate from the recruitment of a B chromosome, which are dispensable chromosomes that are found variably in populations and species (Yoshida et al. 2011).

Moths and butterflies commonly have a ZW sex chromosome system in which females are heterogametic and show a lack of recombination (Turner & Sheppard 1975). Whilst the Z chromosome has been shown to be highly conserved across the Lepidoptera (Fraïsse et al. 2017), the origin and evolution of the W chromosome remains unclear and several putative origins have been hypothesised. First, some recent evidence has been shown to suggest that the W chromosome originated from the same homologous chromosome pair as the Z (Dai et al. 2022)(Figure 1A). However, the possible absence of W chromosomes in early-diverging ditrysian lineages (a lepidopteran clade containing all butterflies and most moths) and the deep conservation of the Z chromosome have been suggested to be evidence of an ancestral Z0 sex determination system for Lepidoptera, and an origin for the W chromosome that is independent of the Z chromosome (Dalíková et al. 2017; Fraïsse et al. 2017; Hejníčková et al. 2019; Lukhtanov 2000; Sahara et al. 2012). Two main alternative hypotheses have been proposed for the origin of the W chromosome (that is independent of the Z chromosome): evolution from a B chromosome (Dalíková et al. 2017; Lewis et al. 2021; Lukhtanov 2000) (Figure 1B), or evolution from the homologous pair of an autosome that fused to the Z chromosome (Sahara et al. 2012)(Figure 1C). Furthermore, and irrespective of the specific origin, the number of evolutionary events leading to the formation of the W chromosome has also been debated. That is whether all W chromosomes in the Lepidoptera share a common evolutionary origin of if their inception was the result of multiple independent events, with the most recent evidence supporting the latter (Dai et al. 2022; Lewis et al. 2021)(Figure 1D-E).

**Fig. 1.**
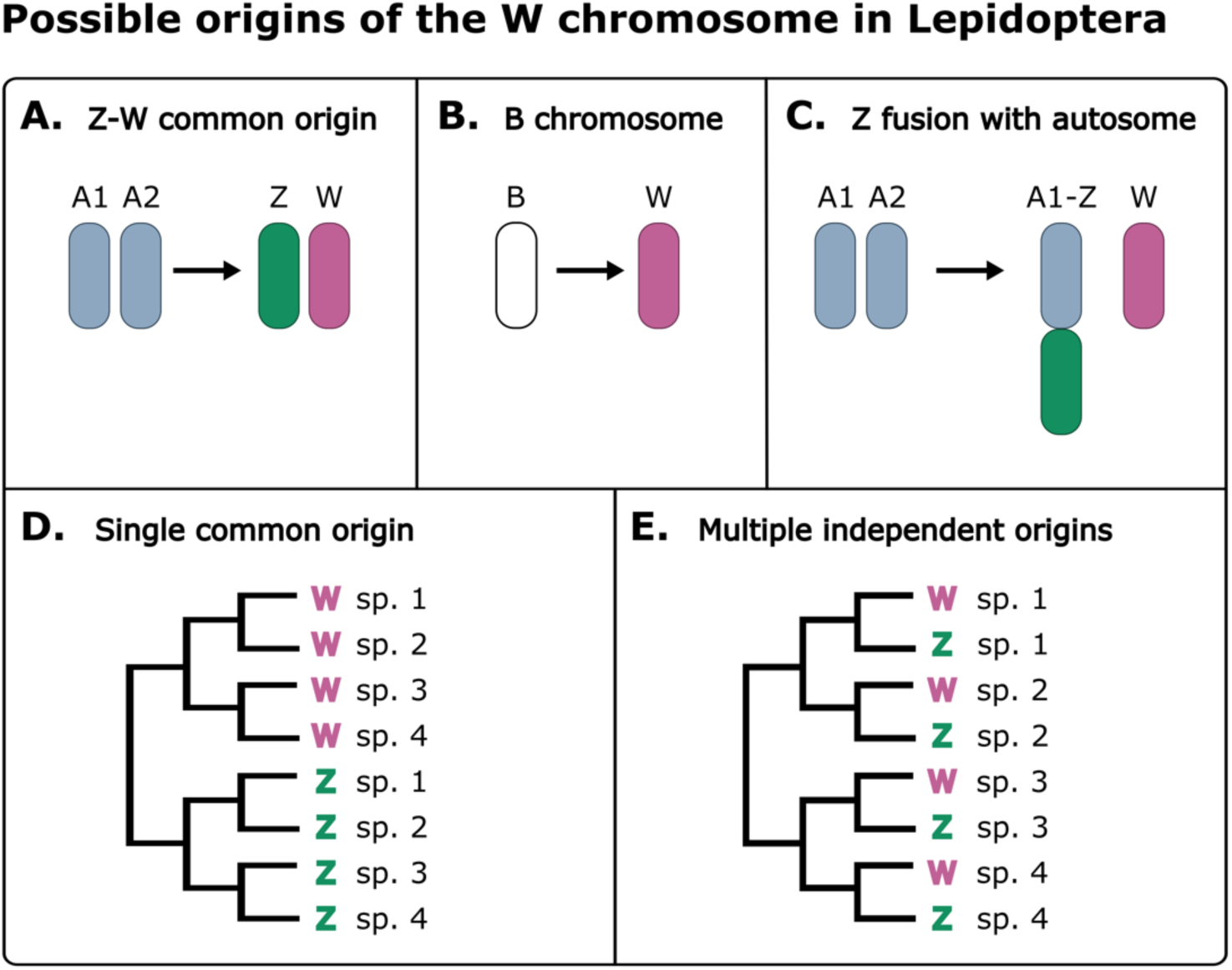
Possible origins of the W chromosome in Lepidoptera. **A.** One hypothesis on the origin of W chromosomes is that Z and W evolved from the same autosomal pair. **B.** Another hypothesis is that the W chromosome evolved from a B chromosome, which are non-essential chromosomes that vary in number in populations and species. **C.** Finally, the inception of W chromosomes could have involved the fusion of the Z chromosome to an autosome and subsequent formation of the W from the autosomal pair. **D.** In any of these cases, if the W chromosome has originated only once in Lepidoptera, W chromosomes of all species would be more similar to each other than to the Z chromosomes. **E.** Whereas if they evolved multiple times from the same autosomal pair as the Z, W chromosomes would be more similar to the Z of the same species than to other Ws. While, multiple B chromosome recruitments would result in a lack of similarity between Z and W chromosomes as well as among Ws.

Despite interest in understanding the evolution of the W chromosome in the Lepidoptera, the absence of high-quality reference genomes containing assembled W chromosomes has limited its study. The elevated repeat and TE content have made its assembly challenging until recently. Long read sequencing technologies and the decrease in sequencing price have enabled the production of high-quality Lepidopteran genome assemblies containing W chromosomes, which makes it increasingly possible to elucidate the enigmatic origin of the Lepidoptera W chromosome.

*Hypolimas* butterflies, commonly known as eggflies, are a phenotypically diverse genus that has served as a model for the study of ecology and evolutionary biology. Many *Hypolimnas* species are mimics of toxic species, which has shaped the diversification of wing colour pattern in the genus (Vane-Wright et al. 1977). Historically, two species, *Hypolimnas bolina* and *Hypolimnas misippus*, have received most of the attention. *Hypolimnas bolina* and *H. misippus* diverged 8 million years ago and share many similarities; both have a pantropical distribution and the females of both species are polymorphic Batesian mimics of toxic models (Marsh et al. 1977; Sahoo et al. 2018; Smith 1976). In contrast, males are monomorphic and have retained what is likely to be the ancestral phenotype of white-spotted black wings. Nonetheless, the two species also differ in many aspects. *Hypolimnas misippus* females are mimics of the four morphs of the African Queen, *Danaus chrysippus*, while *H. bolina* females are mimics of several *Euploea* species. *Hypolimnas bolina* has also received special interest for its coevolution with the endosymbiont *Wolbachia* (Charlat et al. 2009; Dyson et al. 2002). In *H. bolina*, *Wolbachia* has a male-killing phenotype that promotes spread of the endosymbiont through females but has counter-evolved a suppressor locus that rescues male butterflies (Hornett et al. 2006).

The diversity in phenotype, precision of mimicry and intricate coevolution, make *Hypolimnas* a remarkable genus for evolutionary biology studies. However, there are few genomic resources to date. Here, we present chromosome-level assemblies for *H. misippus* (HypMis_v2) and *H. bolina* (HypBol_v1) and use our high-quality *H. misippus* assembly containing the Z and W chromosomes to explore the evolution of sex chromosomes in Lepidoptera. First, we compare synteny and TE content between the two *Hypolimnas* assemblies. Next, we evaluate the annotation completeness and gene content by comparison to the closely related painted lady butterfly, *Vanessa cardui.* Then, we investigate the hypothesis of a B chromosome origin of the W chromosome by comparing homology between the W chromosome, autosomes, and Z chromosome within *H. misippus*. Finally, we examine the origin of the W and Z chromosomes across the Lepidoptera by analysing synteny between *Hypolimnas misippus* and a diverse set of 10 Lepidoptera species in different ditrysian families.

## Results

### Genome assemblies and synteny between HypMis_v2 and HypBol_v1

The size of the final assemblies were 438.07 MB for HypMis_v2 and 444.68 MB for HypBol_v1, assembled into 218 and 59 scaffolds respectively. HypMis_v2 was sequenced using a trio-binning strategy, which, by using a combination of short and long-read sequencing of the parents and offspring of a cross, allows for the assembly of two parental haplotypes (Yen et al. 2020). We assembled both parental haplotypes to a quasi-chromosome level and then scaffolded haplotype 2, which corresponds to the maternal haplotype, using Hi-C data. Hereafter, all mentions of the *H. misippus* assembly, HypMis_v2, refer to the Hi-C scaffolded haplotype 2 assembly.

HypMis_v2 was assembled into 32 chromosome-level scaffolds (>6 MB; 30 autosomes and the Z and W chromosomes) and 186 unplaced scaffolds smaller than 1MB (Figure 2B) and has an N50 of 14.6 MB; while HypBol_v1 was assembled into 59 scaffolds all placed onto 31 chromosomes, with an N50 of 15.2 MB. Thus, both species have retained the ancestral karyotype of 31 chromosomes. This karyotype is present in other Nymphalinae species such as the painted lady *Vanessa cardui*, which has a comparable genome size (424.8 Mb) to the two *Hypolimnas* species. In general, all chromosomes were slightly larger in HypBol_v1 compared to HypMis_v2 with only 4 exceptions (chromosomes 13, 14, 28 and 31; Figure 2C). When aligning the two *Hypolimnas* assemblies, there were no fusions or fissions among chromosomes, but multiple large rearrangements were observed (Figure 2A; Supplementary Figure 1). The only caveat to these conclusions is that HypMis_v2 was used to scaffold HypBol_v1 before using the linkage map, which could theoretically have caused us to miss differences in the *H. bolina* chromosomal structure. However, the linkage map had 31 linkage groups, providing an independent line of evidence that the two species have the same karyotype. Moreover, there were no conflicts between the linkage map and the assembly curated with ragout, and therefore no evidence of inter-chromosomal rearrangements (e.g. translocations) between the species. Finally, 12 large inversions on multiple chromosomes are apparent in the alignment of the two assemblies (Supplementary Figure 1).

**Fig. 2.**
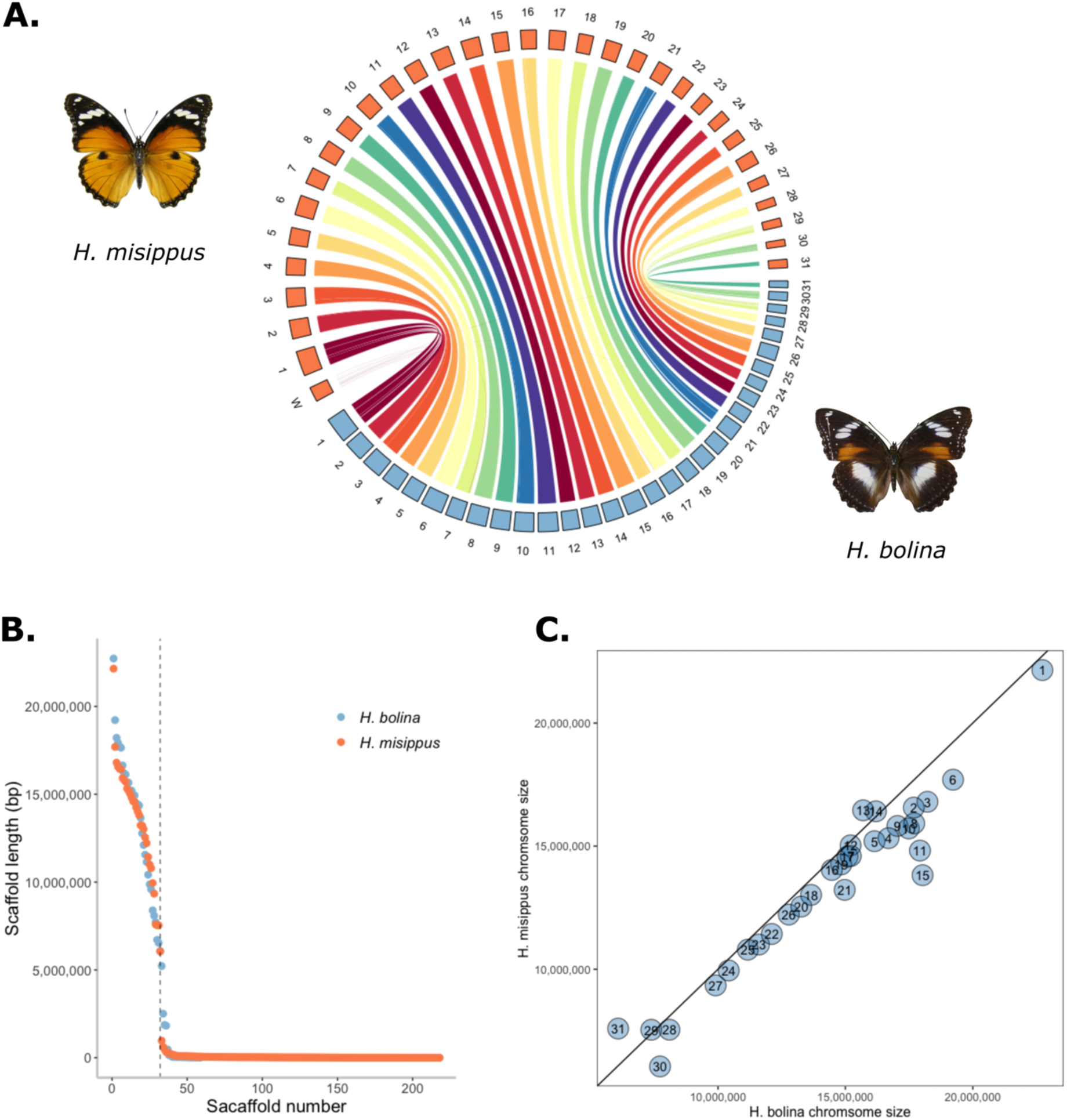
Chromosome level assemblies for *Hypolimnas misippus* and *H. bolina*. **A.** Chromosomal synteny is conserved in both species. **B.** Chromosome level scaffolds have been assembled for both species. **C.** In general, *H. bolina* chromosomes are larger. Chromosome 1 refers to the Z chromosome. *H. bolina* image modified from (MCZ Harvard University 2023) under CC BY-NC-SA 3.0.

### Transposable elements and repeat content

In total, the two *Hypolimnas* genomes contain similar levels of repeats, 43.86% for HypBol_v1 and 42.34% for HypMis_v2, which are slightly higher than in the painted lady *Vanessa cardui* (37.27%) of comparable genome size (424.8 Mb). The composition of repeats is similar in the two *Hypolimnas* genomes, while they differ substantially from the painted lady (Figure 3). The contribution of rolling-circles (also known as *Helitrons*) and DNA-transposons is more than double in *Hypolimnas* than in the painted lady, while retroelements are >1.5 fold higher in the painted lady (Figure 3). This suggests a shift in TE activity, with *Helitrons* and DNA-transposons playing a more important role in *Hypolimnas* species. In both *Hypolimnas*, there has been both a recent and a more ancient expansion of *Helitron* family transposable elements (Figure 3B). Finally, the percentage of unclassified repeats and GC content is broadly the same in the three species.

**Fig. 3.**
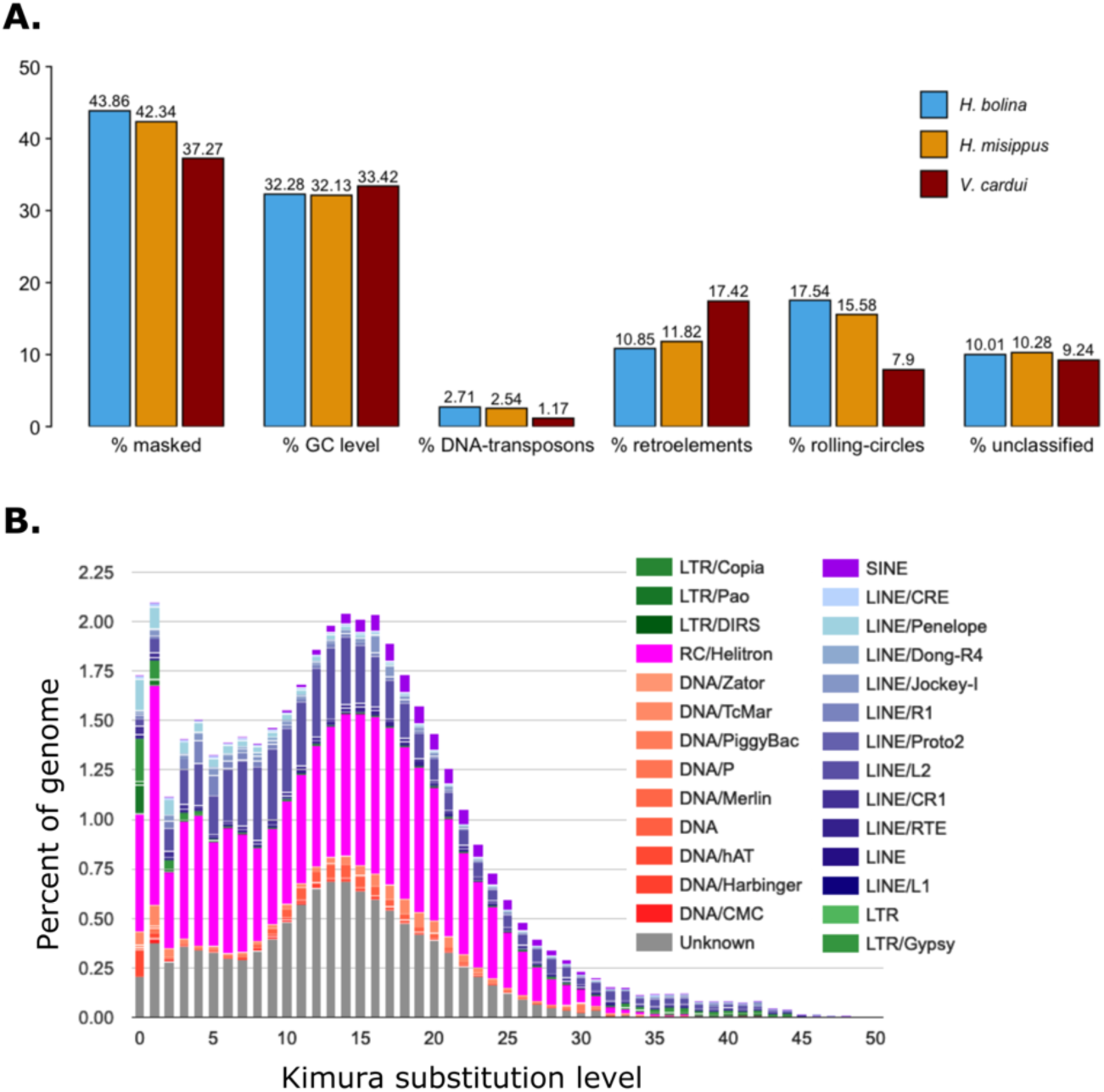
**A.** Repeat content of the *Hypolimnas bolina, Hypolimnas misippus* and *Vanessa cardui* assemblies. **B**. Repeat landscape of the *H. misippus* assembly. Helitrons and LTR (Long Terminal Repeat) retrotransposons have undergone a recent expansion, as seen in the higher percentage of the genome covered by *Helitrons*.

### Gene content and completeness of the annotation

In total 19,721 genes were annotated in HypBol_v1 and 21,784 coding mRNAs including all isoforms of the same gene; while for HypMis_v2 there were 20,293 genes and 22,468 coding mRNAs were predicted (Supplementary Table 1). These numbers are higher than other Nymphalidae species such as the painted lady butterfly (*Vanessa cardui)*, whose latest annotation (v2.1) includes 13,223 genes and 19,836 mRNAs, with the number of genes being considerably larger in the two *Hypolimnas* species (Supplementary Figure 2, Supplementary Table 1). Analysis with BUSCO using the *insecta_odb10* benchmarking set showed that the completeness of the genome and annotation were 98.7% and 98.5% for HypMis_v2, and 94% and 92.4% for HypBol_v1 (Table 1). These scores are comparable to other published Nymphalidae assemblies and annotations such as that of *Danaus chrysippus* (Singh et al. 2022) or the small tortoiseshell butterfly, *Aglais urticae* (Bishop et al. 2021), with those for *HypBol_v1* slightly lower possibly due to errors introduced by Nanopore sequencing, which has a high per base error rate.

**Table 1.**
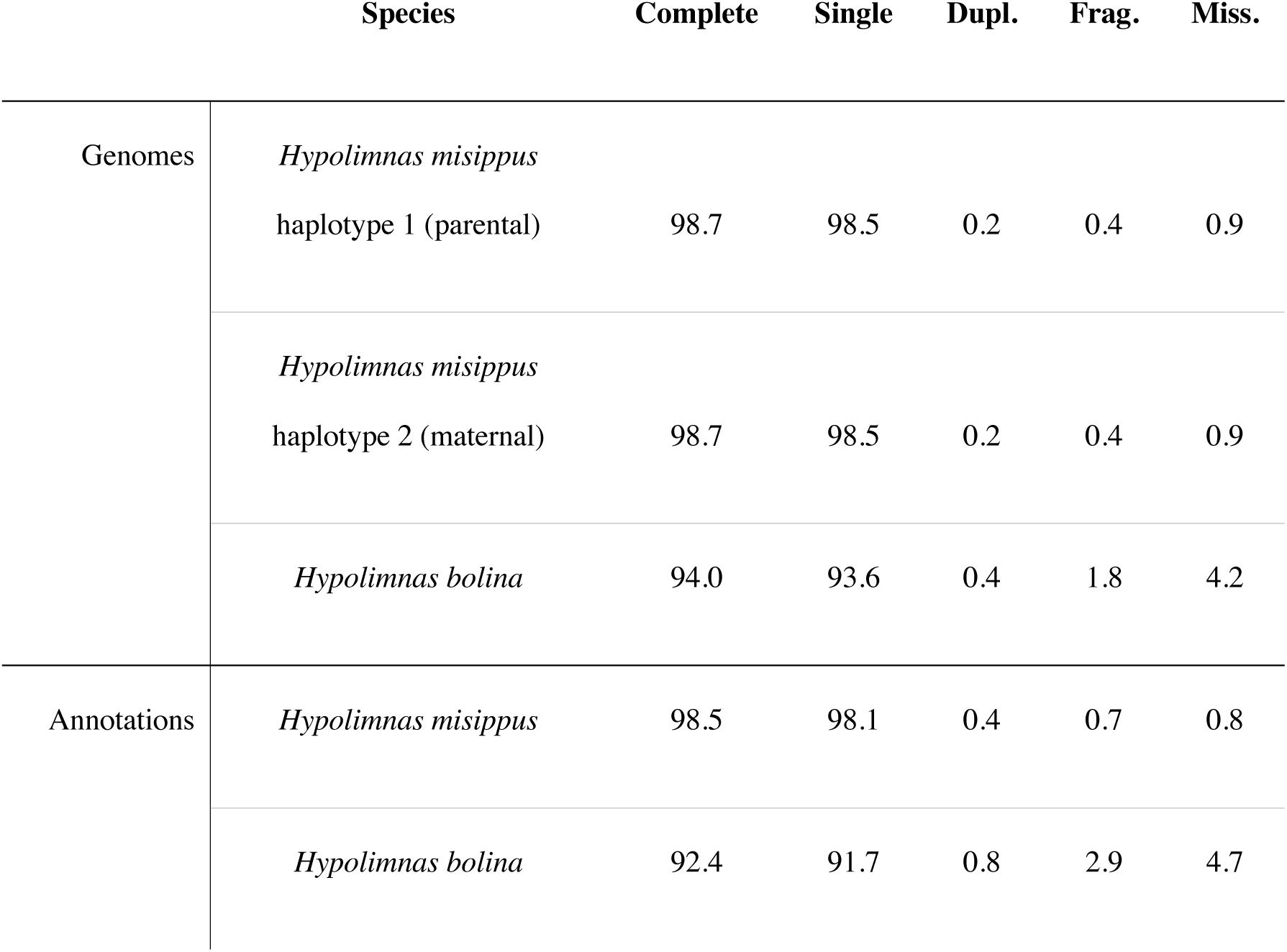
BUSCO scores for the genome assemblies and gene annotations of *H. misippus* and *HypBol_v1* calculated using the Insecta_odb10 (n=1367) set of genes.

Despite having a larger number of genes and mRNAs, the two *Hypolimnas* annotations show a smaller total number of exons and introns than *V. cardui* (Supplementary Figure 2). This is because the two *Hypolimnas* annotations have more single exon genes and their mRNAs have, on average, a smaller number of exons, which might be a difference produced by the distinct annotation pipelines. The total length of mRNAs and genes in the two *Hypolimnas* is shorter then in *V. cardui*, which results in a smaller percentage of the genome covered by them. However, this trend is different for exons, which are on average the same length in the three species, have a comparable total length and cover a similar percentage of the genome. Thus, the longer total length of mRNAs and genes in *V. cardui* is due to a total longer length of introns, due to a higher number of introns per mRNA and an average longer length.

### Mixed evidence for the origin of W chromosomes from the same autosome pair as the Z

Using our HypMis_v2 assembly that contains a W chromosome, we set out to explore the evolutionary origins of the W chromosome in Lepidoptera. First, we wanted to clarify whether the W chromosome had originated from a B chromosome or form the same autosomal pair as the Z (Figure 1A-B). To specifically test the hypothesis that the Z and W have a common autosomal origin, we used an approach that searches for syntenic blocks to test for sequence homology between the *H. misippus* W chromosome and the remaining 31 chromosomes. We would expect that if the W and the Z share a common autosomal origin, they might share tracts of homologous sequence that date back to their common ancestor. As such, we might expect that the Z and W would each share more syntenic blocks with each other than with any of the autosomes. We found that the Z chromosome indeed shares a higher proportion of non-overlapping syntenic blocks with the W (Figure 4A). However, the W chromosome is the chromosome with the highest percentage of repeat regions (71% in *H. misippus*), which could interfere with the analysis (i.e. if the sex chromosomes share many common repetitive sequences that may have accumulated independently on each sex chromosome). Furthermore, the Z chromosome is the longest chromosome in *H. misippus* and has the highest absolute length of repeats (Supplementary Figure 3 and 4). To take this into account, we excluded all matches found in repetitive regions. With this correction, the Z chromosome shared the second highest proportion of syntenic blocks with the W after chromosome 2.

**Fig. 4.**
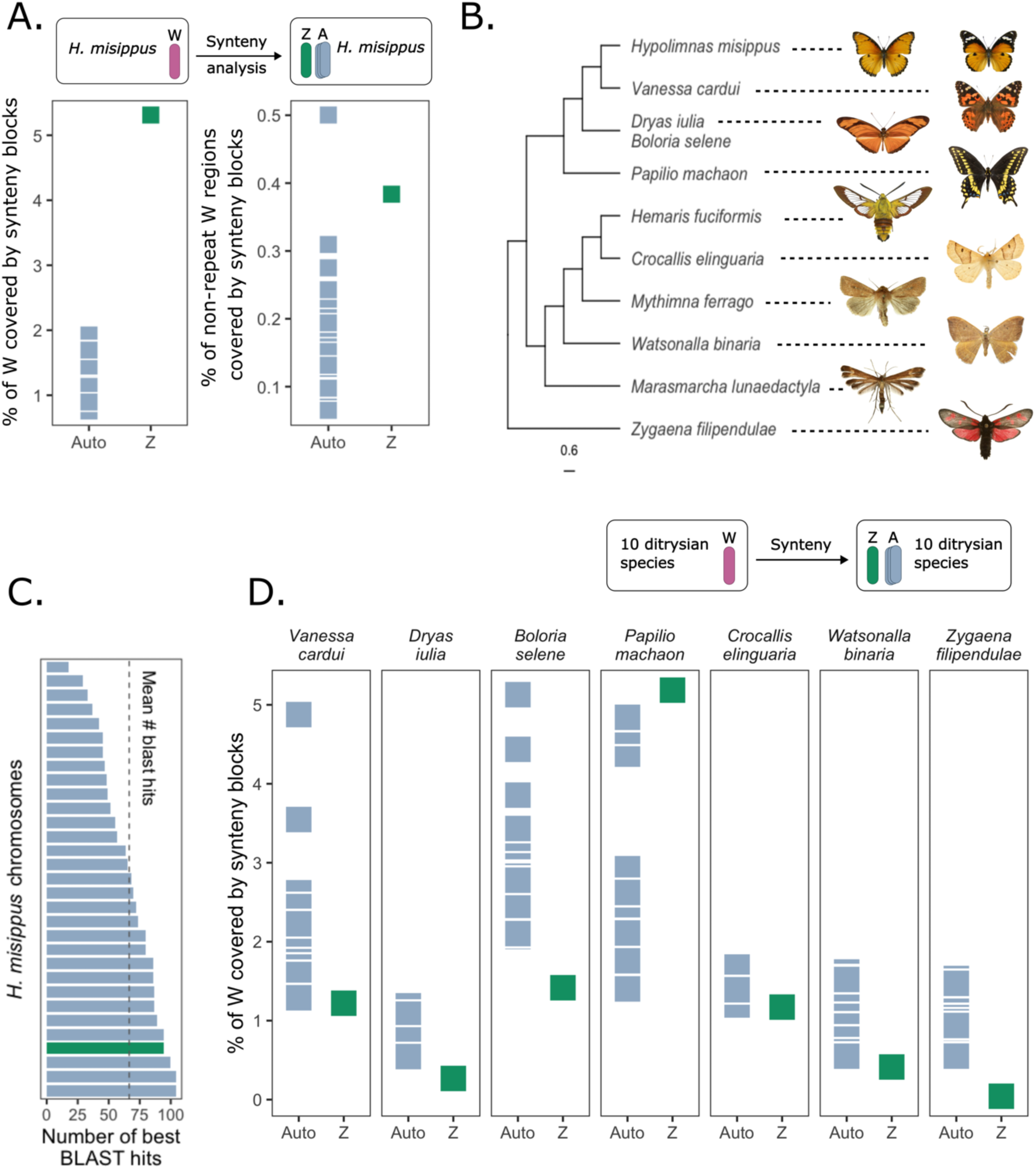
Genomic synteny of the *H. misippus* W chromosome provides mixed evidence on its origin. **A.** Synteny analysis of the *H. misippus* W chromosome and the autosomes and Z chromosome shows that the W and Z chromosome have the highest similarity. The analysis of non-repeat regions shows that the Z chromosome is the second most similar chromosome to the W after chromosome 19 (right). A schematic of the method is shown on top. **B.** Phylogenetic tree modified from Kawahara, et al 2019 (Kawahara et al. 2019) including the species used in the synteny analysis. **C.** Number of best BLAST hits of the proteins found in the *H. misippus* W chromosome to the other chromosomes. Numbers summarised by chromosome with the Z shown in green. **D.** Synteny analysis of the W chromosomes of several Lepidoptera genome assemblies compared to the autosomes and Z of the same species. A schematic of the method is shown on top. *Hemaris fuciformis* image modified from Didier Descouens under CC BY-SA 4.0. *Dryas iulia* image modified from (Moore 2023) under CC-BY. *Watsonalla binaria* image modified from (Lennuk 2023) under CC BY-SA 4.0.

To further explore the possible homology between the W and Z chromosomes, we extracted the predicted amino acid sequences of all 1,394 annotated proteins in the *H. misippus* W chromosome and searched for homology in the remaining chromosomes using BLASTp. In contrast to the synteny results, we found that the Z chromosome placed 4^th^ in the number of best BLAST protein hits with 94 proteins mapping to it, while 104 mapped to chromosome 2, 3 and 14 (Figure 4C). The average number of best BLAST hits was 66. Interestingly, chromosome 2 showed the highest sequence similarity in non-repeat regions and the highest number of best BLAST hits (Figure 4C).

In view of the differences between these results and previously published homology tests in other species, we decided to expand the analysis to include other Lepidoptera species. We searched for syntenic blocks between the W and the other 31 chromosomes in 6 diverse taxa: *Dryas iulia, Boloria selene, Papilio machaon, Crocallis elinguaria, Watsonalla binaria,* and *Zygaena filipendulae*. These species were chosen as they have a publicly available high-quality assembly containing a W chromosome (Supplementary Figure 5) and cover distinct ditrysian lineages (including 4 superfamilies: *Zyganoidea, Papilonoidea, Drepanoidea* and *Geometroidea*). A low percentage of the W chromosome was covered by synteny blocks when comparing it to the Z in any species (suggesting no homology) except for *P. machaon*, which contrasts with the results in *H. misippus* (Figure 4D). However, these analyses were performed using unmasked genomes and including all regions, which could be interfering with the results.

### Comparisons of W chromosomes reveals a possible single common origin in ditrysian lineages

The second question we wanted to clarify was whether the W has a single origin in the Lepidoptera or evolved independently multiple times (Figure 1D-E). To explore the evolution of W chromosomes within ditrysian lineages, we compared the *H. misippus* W chromosome to a diverse set of 10 Lepidoptera species, including the 6 species from the previous analysis plus *Vanessa cardui, Hemaris fuciformis, Mythimna farrago* and *Marasmarcha lunaedactyla.* These species cover the breadth of ditrysian lineages, including 8 distinct superfamilies: *Zyganoidea, Papilonoidea, Pterophonoidea, Pyraloidea, Drepanoidea, Noctuoidea, Geometroidea* and *Bombycoidea*. We searched for syntenic blocks between the *H. misippus* W chromosome and all the chromosomes of each target species (including the W; Figure 5A). This allowed us to test whether W chromosomes are more similar to each other than to any autosome or the Z chromosome, which would provide evidence of a common origin. When comparing the *H. misippus* W chromosome to these species, we found that there is a high proportion of syntenic blocks among W chromosomes (Figure 5A). The highest degree of sharing is seen between W chromosomes of closely related species such as *H. misippus* and *Vanessa cardui* (45 MYa), but is also true even for highly divergent lineages such as *Zygaena filipendulae,* which diverged from *H. misippus* 156 MYa (Kawahara et al. 2019). Interestingly, there is no consistent pattern in the distribution of syntenic blocks along the *H. misippus* W chromosome (Supplementary Figure 6), which might suggest that different regions show homology with the *H. misippus* W chromosome across species and chromosome types (W, Z, and autosomes).

**Fig. 5.**
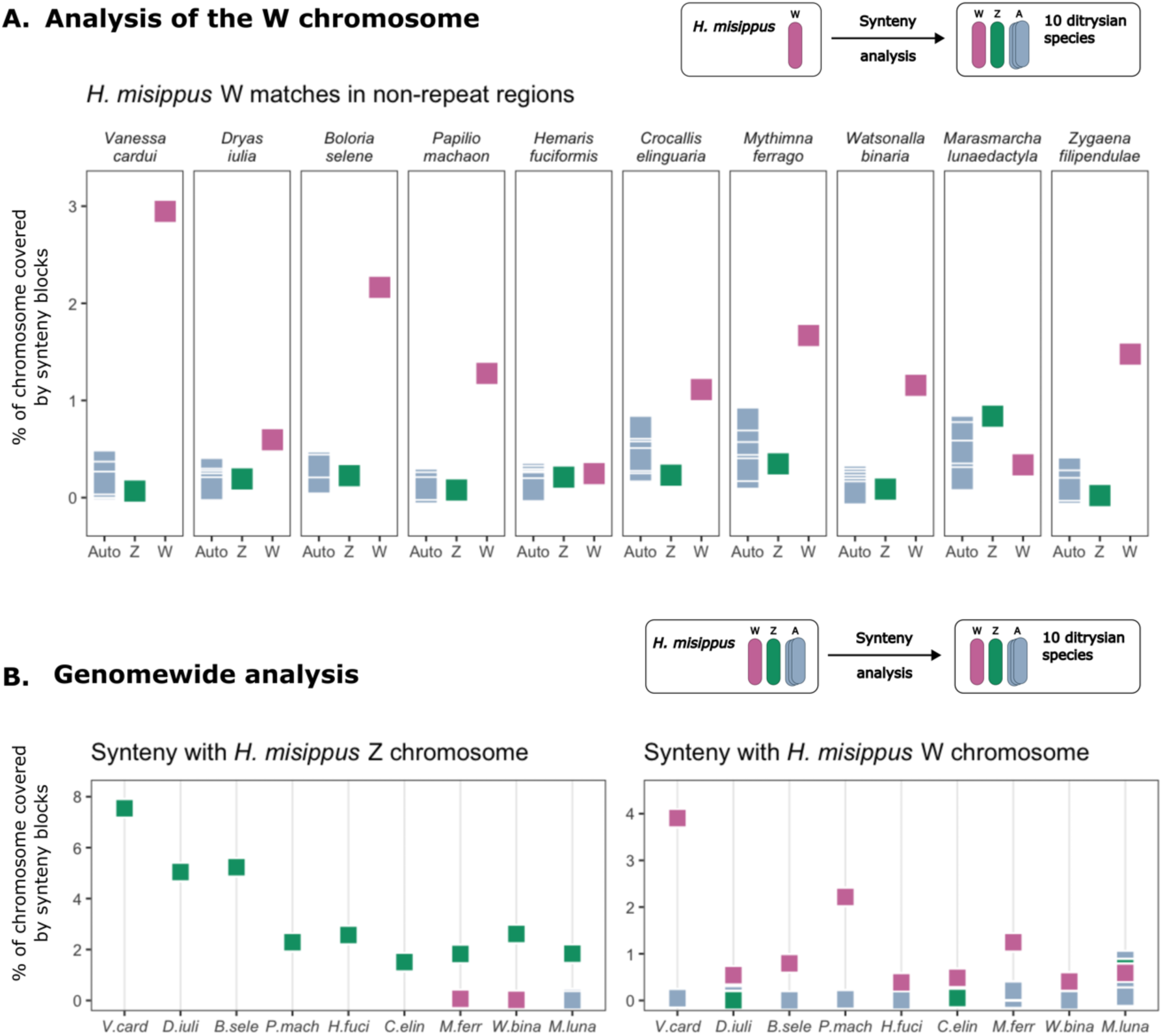
Genome wide comparisons suggests a single origin of ditrysian W chromosomes and shows deep conservation of the Z chromosome. **A**. Percentage of query *H. misippus* W chromosome covered by synteny blocks of the 32 chromosomes of 10 ditrysian species. W *H. misippus* chromosome used as query (a schematic of the method is shown on top). **B.** Subset results of genome wide analysis. Percentage of Z (left) and W (right) chromosome covered by synteny blocks to other ditrysian species. W chromosome shown in pink, Z in green and autosomes in grey. A schematic of the method is shown on top. Species names have been abbreviated: *Vanessa cardui, Dryas iulia, Boloria selene, Papilio machaon, Hemaris fuciformis, Crocallis elinguaria, Mythimna 60arrago, Watsonalla binaria, Marasmarcha lunaedactyla* and *Zygaena filipendulae*.

We then performed the same analysis including all chromosomes of *H. misippus* as the query. With this we could test for homology between all autosomes and sex chromsomes of *H. misippus* with each chromosome of the target species. Unlike the above method, this approach produces only primary matches for each query chromosome and is thus efficient at finding homologous pairs. This approach yielded the same results as the former, in which the W chromosome of *H. misippus* shares the greatest proportion of syntenic blocks with the W of most species, with *Marasmarcha lunaedactyla* being the one exception (Figure 5B right). Finally, the percentage of the Z chromosome covered by synteny blocks was the highest with other Z chromosomes in all species comparisons (5B left), suggesting that it is highly conserved.

### A multi-species comparison reinforces the hypothesis of the single origin of the W chromosome and reveals a neo-Z chromosome

In light of the above results suggesting a single origin of the W chromosome in ditrysian lineages, we decided to explore this further by comparing the analysed ditrysian assemblies with each other. We performed pairwise searches for syntenic blocks between species pairs and observed consistent results. First, the Z chromosome is deeply conserved in multiple species pairs and the proportion of syntenic blocks between Z chromosomes of different species decays with species divergence in a similar manner to autosomes (Supplementary Figure 7 and 8). Second, the W chromosome shows a more variable pattern of sequence sharing between species but shows evidence for conservation even between distant species pairs (Supplementary Figure 8). Nonetheless, several of the comparisons found no syntenic blocks between W chromosomes, again indicating that the syntenic blocks shared with the *H. misippus* W differed between target species.

Finally, the cross-species comparison revealed that the Z chromosome of *Marasmarcha lunaedactyla* (chromosome name OV181339.1; identity as the Z chromosome assigned in the public assembly) presented high levels of synteny with chromosome 19 of most other species (Supplementary figures 9 and 10). This could suggest a historical fusion event between the ancestral chromosome 19 and the Z chromosome of *M. lunaedactyla* creating a neo-Z chromosome. The pattern of syntenic blocks seen in the OV181339.1 chromosome was similar for all species compared, with approximately half of the chromosome showing homology to chromosome 19 and half to the Z chromosome (Supplementary Figure 10). Furthermore, the pattern of synetny with other species showed that translocations between the parts belonging to the ancestral Z and ancestral chromosome 19 have resulted in an interleaved pattern of homology. Consistent with these results, BUSCO matches of chromosome OV181339.1 are shared with chromosome 19 and chromosome 1 (the Z chromosome) of *Melitaea cinxia* (Supplementary Figures 9 and 10).

## Discussion

*Hypolimnas* species have become a focus for studies of evolutionary genetics, due to their mimetic colouration and co-evolution with *Wolbachia* parasites. Here, we have assembled chromosome level reference genomes for *H. misippus* and *H. bolina* and RNA-informed annotations for both genomes. HypBol_v1 represents the first assembly available for *Hypolimnas bolina,* while HypMis_v2 is a significant improvement on a previously published genome, with higher contiguity and higher N50 and BUSCO scores. Furthermore, the annotation is also improved as we used RNA-seq data from *H. misippus* to inform our annotation, as opposed to only homology methods. We present these two useful resources and use the *H. misippus* assembly to shed light on the evolutionary origins of W chromosomes in Lepidoptera.

Several models have been proposed to explain the origin and evolution of the W chromosome in Lepidoptera (Dai et al. 2022; Fraïsse et al. 2017; Lukhtanov 2000; Sahara et al. 2012; Berner et al. 2023). The deep conservation of the Z chromosome and its apparent lack of homology with the W have been suggested to be evidence of a B chromosome origin of the W chromosome (Fraïsse et al. 2017). Outside the Lepidoptera, it has been commonly found that heteromorphic sex chromosomes retain a degree of similarity; evidence of their shared autosomal origin (Wright et al. 2016). Thus, if the W chromosome originated from the same homologous autosome pair as the Z, we might expect to find some residual sequence similarity between the W and Z, as some recent evidence seems to suggest. However, opposing evidence has also been presented, suggesting a lack of similarity between W and Z chromosomes which could point to the evolution of the W from a B chromosome (Fraïsse et al. 2017; Lewis et al. 2021). Furthermore, the lack of similarity among W chromosomes of Lepidoptera species has led to the suggestion of multiple independent recruitments from B chromosomes (Dai et al. 2022; Lewis et al. 2021). Taking this into account, we set out to address two questions. First, did the W chromosome evolve from the same homologous pair of autosomes as the Z or from a B chromosome? Second, do W chromosomes in Lepidoptera share a common origin, that is, did they evolve once or multiple times independently? Our assembly of the W chromosome of HypMis_v2 is an ideal system to explore these questions, as its large size (13.23 Mbp) compared to other W assemblies (e.g. 1.87 Mbp in *Zygaena filipendulae* or 2,1 in *Dryas iulia*) increases our chances of detecting homology even in scenarios of extensive degradation.

Using tests for homology between the W and remaining 31 chromosomes in *H. misippus* and other Lepidoptera, we show that there is some evidence for homology between the W and the Z chromosome, suggesting a possible common origin. These results differ from the homology tests performed in *Danaus plexipuss* (Lewis et al. 2021), *Kallima inachus* (Yang et al. 2020) and *Dryas iulia* (Lewis et al. 2021), where the evidence was inconsistent with a common origin of the W and the Z chromosome. Instead, our findings are more in line with results in the Asian corn borer, *Ostrinia furnacalis,* a moth belonging to the Pyraloidea, and *Pieris mannii* which also suggest a common autosomal origin of the Z and the W chromosome (Dai et al. 2022; Berner et al. 2023). Contrastingly, our results based on protein-coding genes contrast with the analysis of overall homology and suggest that the W chromosome did not evolve from the same homologous autosomal pair as the Z. Nonetheless, as a result of the arrest of recombination and subsequent accumulation of mutations, sex specific chromosomes such as the W are often gene poor and highly degenerated compared to their autosomal counterparts (Wright et al. 2016). Additionally, lepidopteran W chromosomes have been shown to evolve rapidly (Yoshido et al. 2013; Vítková et al. 2007). Thus, the lack of homology between proteins of the W and other chromosomes is expected given the rapid turnover seen in the W chromosome and is in line with results shown in other Lepidoptera species (Yang et al. 2020; Singh et al. 2022; Lewis et al. 2021). Similarly to the protein-coding analysis, the low proportion of synteny blocks between the W and the Z in five of the seven species studied could suggest a B chromosome origin or could be the result of the rapid evolution and degradation of the W, which could impede the detection of ancient homology (Yoshido et al. 2013; Vítková et al. 2007). Taken together, the homology tests between the W and remaining 31 chromosomes in *H. misippus* and other Lepidoptera provide mixed evidence for the origin of the W from the same homologous pair of autosomes as the Z or from a B chromosome. Alternatively, our finding of synteny blocks between the *H. misippus* W and the Z could point to a fusion of the Z chromosome with an autosome and subsequent formation of the W (Figure 1C). This hypothesis would fit the mounting evidence supporting a Z0 system in non-ditrysian Lepidoptera and early diverging ditrysian lineages (Fraïsse et al. 2017; Hejníčková et al. 2019; Lukhtanov 2000; Sahara et al. 2012).

The second question we wanted to address was whether the W has a single origin in the Lepidoptera or evolved independently multiple times. Previously, the lack of similarity among W chromosomes of Lepidoptera species has been suggested to be the result of multiple independent recruitments from B chromosomes (Lewis et al. 2021). Similarly, phylogenies of W genes in the Asian corn borer could suggest multiple independent origins of the W chromosome from the same autosomal pair as the Z (Dai et al. 2022). Regardless of the specific event that led to the formation of the W chromosome (i.e., from a B chromosome or from an autosome), if the W chromosomes of ditrysian lineages share a common origin, we would expect them to share homologous sequence tracts that date back to their common ancestral chromosome, and therefore we expected to find more syntenic blocks between W chromosomes than between W and and Z or W and autosomes. Comparing the *H. misippus* W chromosome to the genomes of 10 other ditrysian species, our findings matched this expectation in all but one species. These results suggest that W chromosomes of ditrysian lineages may share a single common origin, which contrasts with evidence of rapid turnover of W chromosomes in Lepidoptera (Dai et al. 2022; Lewis et al. 2021). However, it is possible that some species have retained the ancestral W, while some other have recruited other B chromosomes as suggested. For example, in our test for homology between the *H. misippus* W chromosome and each of the chromosomes of *Marasmarcha lunaedactyla*, the W chromosome of *M. lunaedactyla* did not stand out above the Z and some autosomes. Our results also show unambiguous evidence for homology among Z chromosomes of the studied ditrysian species, which is consistent with evidence suggesting deep conservation of the Z chromosome across the Lepidoptera (Fraïsse et al. 2017).

Overall, we present evidence of a single common origin of W chromosomes in the Lepidoptera. Furthermore, our results hint at a possible origin from the same autosomal pair as the Z or perhaps to a Z-autosome fusion event that culminated with the formation of the W. Crucially, the rapid evolution of the W chromosome could cause the lack of similarity observed between Z and W chromosomes in some species and hinders the elucidation of its origin. Further studies including more species across the Lepidoptera are necessary to clarify whether the W chromosome evolved from an autosome or from a B chromosome, and to shed light on the possible shared origin of W chromosomes, particularly including non-ditrysian lineages.

## Materials and methods

### *Hypolimnas misippus* butterfly rearing and cross preparation

A trio-binning approach was used for sequencing of the *Hypolimnas misippus* assembly, which consists of sequencing the parents and offspring from one family. First, butterflies were obtained from Stratford-upon-Avon Butterfly Farm, UK, and reared in greenhouses in Madingley, Cambridge, UK. Larvae were fed *Portulaca oleracea* and *Portulaca quadrifida*. Adult butterflies were kept in a large cage (1.5 meters x 1.5 meters x 2 meters) and observed until mating occurred when mating pairs were transferred to separate smaller cages during copulation. One mated pair was used for trio-based genome sequencing, and their offspring were reared until pupation and then flash frozen as pupae in liquid nitrogen.

### *H. misippus* Trio binning genome assembly

One *H. misippus* family was used for sequencing and trio-binning genome assembly. DNA was extracted using MagAttract HMW DNA Kit (QIAGEN). The two parents were sequenced using Illumina short-read sequencing, which resulted in 368.67 and 346.56 million read pairs and a total yield of 55.67 and 52.33 Gbp of data from the mother and father respectively. One of the offspring was sequenced using PacBio long-read sequencing (total yield of 13.46 Gb of data and N50 of 13,493). Trio binning enables the independent assembly of the two parental haplotypes. First, yak-r55 (Li 2022) was used to create a kmer database from the parental Illumina data. Then, hifiasm-0.7-r256 (Cheng et al. 2020) was run in trio mode to assemble the HiFi PacBio long-read data, followed by the purging of haplotypes and overlaps using purgedups v1.2.3 (Guan 2022). Haplotype 1 corresponds to the paternal haplotype, while haplotype 2 corresponds to the maternal haplotype. All subsequent analysis were performed using haplotype 2.

### Curation of HypMis_v2 with HiC

To place the assembled scaffolds into chromosomes, Hi-C sequencing was used, which is a chromosome conformation capture technology that provides information about the 3-D interactions between genomic loci. An offspring from an unrelated mating was flash frozen in liquid nitrogen and used for Hi-C sequencing and analysis. 386.99 million read pairs were produced, which represented 58,45 Gb of Hi-C data. Fastq files were converted to cram (two files, one for each read pair end), mapped to haplotype 2 (the maternal haplotype). The *H. misippus* assembly was produced after the purgedups stage and processed with Juicer v1.6 (Durand, Shamim, et al. 2016). Juicer transforms raw Hi-C data into a list of contacts, which defines pairs of genomic positions that were in close physical contact during the experiment. Then, the main reference assembly was curated using the 3D-DNA pipeline, which corrects misassembles, anchors, orders and orients fragments of DNA based on the Hi-C data (Dudchenko et al. 2017; Durand, Robinson, et al. 2016). 3D-DNA generated assembly heatmaps as part of its workflow, which indicate the frequency of contact between pairs of genomic locations. Obvious errors in the genome assembly such as large genomic inversions were manually edited by examining the Hi-C heatmaps using the Juicebox tool (Durand, Robinson, et al. 2016; Dudchenko et al. 2018). Finally, the edited assembly was exported as an *.assembly* file and converted to a final fasta assembly file using the ‘run-asm-pipeline-post-review.sh’ script setting –editor-repeat-coverage to 6.

### HypBol_v1 genome curation with Ragout

The *H. bolina* reference assembly (HypBol_v1) was produced using a combination of Nanopore long-read sequencing and a linkage map. Pupae were purchased from the Stratford-upon-Avon butterfly farm, which obtains farmed lineages from South East Asia. One female pupa was used for sequencing. High molecular weight DNA was extracted following a phenol chloroform and glass capillary hook protocol following(Quick 2018). A library was prepared for Nanopore sequencing using the Nanopore Ligation Sequencing Kit (SQK-LSK 109) following a LSK109 bead free library preparation protocol (Tyson 2020). Post library preparation, a total of 1.78µg DNA remained and was sequenced across 2 R9 chemistry Nanopore MinION flow cells, resulting in 7.35 million reads totalling 14.16 Gb. Adapters were removed using Porechop v0.2.4 (Wick 2022) and reads assembled using redbean (Ruan & Li 2020). Assembled contigs were split into bins using MaxBin2 (Wu et al. 2016) and bacterial contigs and reads were removed from the data using blobtools2 (Challis et al. 2020). This produced an assembly with 13,492 contigs and an N50 of 1.4 kbp.

Leveraging the high synteny expected from the two *Hypolimnas* assemblies, Ragout v2.3 (Kolmogorov et al. 2014, 2018) was used to improve the initial *HypBol_v1* assembly using HypMis_v2 as a reference. First, the two genomes were soft-masked using RepeatMasker (Smit et al. 2015), creating the repeat library based on the genome being masked. Then, Cactus (Paten et al. 2011) was used to align both genomes using Python 3.8. The resulting HAL alignment was converted to MAF format using the hal2maf utility from the HAL program (Hickey et al. 2013). Finally, Ragout was run using the MAF alignment between HypBol_v1 and HypMis_v2 as input.

### Rearing of *H. bolina* individuals

A linkage map was used to improve the HypBol_v1 assembly and place the assembled scaffolds into chromosomes. Two families (178303XX and 182703XX) were reared and sequenced. First, female *H. bolina* purchased from Stratford upon Avon Butterfly Farm of SE Asian origin were mated to wild-caught males from Mo’orea (French Polynesia) at the University of California Berkeley Gump Station research facility. Female SE Asian-Mo’orea F1 hybrid offspring were then mated to pure Mo’orea F1 males. The F2 offspring of one of these crosses is family 178303XX. At the same time, male SE Asia-Moorea F1 hybrids were mated to pure SE Asia F1 females. The F2 offspring of one of these crosses is family 182703XX. Butterflies were kept in a large outdoor cage for mating under observation. Any copulating pairs were separated into a small cage. The mated female was then placed in an oviposition cup containing a small plant e.g. *Asystasia sp*. and allowed to lay eggs. Hatched eggs were moved to a rearing box with suitable food plant e.g. Ipomoea sp. and caterpillars reared until pupation. Pupae were moved to individual cups for emergence, adults left to dry for one day and then used for further matings or stored in −80°C freezer.

### Illumina library preparation of *H. bolina* family samples

The offspring of the *H. bolina* families (F2) were processed to extract the DNA and prepare the libraries for Illumina sequencing. DNA extractions were carried out using a custom protocol using PureLink buffers and homemade magnetic beads. Briefly, a small piece of thorax tissue (1/10) is placed in an 8-tube PCR strip. Then, 45 µL of PureLink Digestion buffer and 10 µL of Proteinase K (20mg/mL) are added, and the mix is incubated at 58°C with shaking (500 rpm) for 2-3 hours. Afterwards, 2 µL of RNAseA are added (DNAse free, 10mg/mL) and incubated for 10min at room temperature. Then, 45 µL of PureLink Lysis buffer is added to the mix and incubated at 58°C for 30 minutes with shaking (500 rpm). Following that, a homemade magnetic bead mix is used to extract the DNA from the lysate. First, 37.5 µL of magnetic beads are added together with 75 µL of lysate to a 96-well plate. After mixing, the samples are incubated for 15 minutes at room temperature, then the plate is placed on a magnetic stand for 10 minutes, the supernatant removed, and the beads cleaned with 80% ethanol. After drying out, 50 µL of 10mM Tris (pH=8) are added to elute the sample and incubated at 45°C for 15 minutes without resuspending. Then, the beads are resuspended and incubated for 20 minutes at room temperature. Finally, the plate is placed on the magnetic stand and, after 10 minutes, the supernatant (the DNA) is transferred to a fresh tube.

The F2 were sequenced using a Nextera-based library preparation at intermediate coverage (~11X). A secondary purification using magnetic SpeedBeads™ (Sigma) was performed prior to Nextera-based library preparation. Libraries were prepared following a method based on Nextera DNA Library Prep (Illumina, Inc.) with purified Tn5 transposase (Picelli et al. 2014). PCR extension with an i7-index primer (N701–N783) and the N501 and N502 i5-index primers was performed to barcode the samples. Library purification and size selection was done using the same homemade beads as above. Pooled libraries were sequenced by Novogene Cambridge, UK. Libraries of the parental samples were prepared and sequenced to ~20X coverage by Novogene Cambridge, UK.

### *H. bolina* linkage map construction and anchoring of the genome

A linkage map was produced with Lep-Map3 (Rastas 2017) and then used to improve the *H. bolina* assembly and place the scaffolds onto chromosomes. First, sequences were mapped to the reference genome using bwa-mem (Li 2013). PCR duplicates were marked using the MarkDuplicates from Picard tools. Sorted BAMs were then created using SAMtools (Li et al. 2009) and genotype likelihoods computed. The pedigree of individuals was checked and corrected using IBD (identity-by-descent) with a random subset of 10% of the markers (1,270,024 SNPs) following the IBD pipeline from Lep-Map3. These markers were also used to construct the linkage map. Scaffolds were anchored into chromosomes based on the linkage map using LepAnchor (Rastas 2020).

### HypBol_v1 polishing with Pilon

After anchoring with the linkage map, three iterations of Pilon v1.24 (Walker et al. 2014) in diploid mode were run to correct the draft assembly by correcting bases, filling gaps and fixing mis-assemblies. Samples used for the Pilon correction were CAM035727, CAM035728, CAM035186, CAM035187, CAM035188, CAM035189.

### Repeat annotation

Once the two final assemblies had been produced, they were each assessed for repeat content using a custom repeat library. First, a repeat database was built and the repeats of the two finished assemblies modelled using RepeatModeler v. 2.0.2a. Each custom library was then combined with the Lepidoptera library extracted from Dfam (Storer et al. 2021). This merged library was used to soft mask the genome using RepeatMasker v 4.1.0 with the cut-off score set to 250 and skipping the bacterial insertion element check. The resulting soft masked assemblies were used for gene annotation.

### RNA-seq sample preparation

To assist with genome annotation, RNA-seq data was obtained from 4 *H. misippus* individuals (2 adults and 2 pupae) and 17 *H. bolina* individuals (6 adults and 11 pupae). Butterflies were purchased from Stratford-upon-Avon Butterfly Farm and kept at room temperature until dissection. 4 tissues were dissected out from *H. misippus* pupae and placed in RNA-later (Sigma): wing discs, thorax, head, and abdomen; while *H. bolina* pupae 3 tissues were dissected: wing discs, thorax-head, and abdomen. Only abdomen, head and thorax samples were dissected from adults of either species. Two pooling strategies were followed: 1) 2 *H. bolina* pupae were pooled by individual, pooling head, thorax, abdomen, and wing discs together and sequenced at high coverage (50M reads). And 2) 4 *H. misippus* and 15 *H. bolina* adults and pupae were dissected into tissues and pooled by species and tissue. Each pooled sample, 4 for *H. misippus* and 3 for *H. bolina,* was sequenced to 20M reads. RNA was extracted using a modified Trizol protocol using the same protocol as in (Brien et al. 2022).

### RNA-seq mapping and gene annotation

First, low quality ends and adaptors from the RNA-seq data were trimmed using TrimGalore! v 0.5.0 (Krueger 2015). Then, the reads were mapped to the soft masked genomes using STAR v 2.5.0a (Dobin et al. 2013). Two rounds of mapping (2pass) were performed, including all the splice junction files in the second round. Then the resulting mapped reads and the soft masked genome assembly were used to generate a gene annotation using BRAKER v 2.1.5 (Brůna et al. 2021), running it a second time to add UTR annotations with options addUTR=on and skipAllTraining (Keilwagen et al. 2019). A single isoform per gene was selected and completeness of the annotation assessed using BUSCO (Seppey et al. 2019; Simão et al. 2015) using the Insecta_odb10 (n=1367) set of genes.

### Homology with Merian units

A growing practice in Lepidoptera genetics is to name chromosomes of reference assemblies by their homology to the ancestral lepidopteran chromosomes, also known as Merian units in honour of the scientist Maria Sibylla Merian (Wulf 2016; Wright et al. 2023) and allow for the tracking of conserved synteny blocks across the phylogeny of the Lepidoptera. To assign homology to the Merian units, lepidopteran_odb10 BUSCO assignment of the *Melitaea cinxia* genome was compared to the lepidopteran_odb10 BUSCO assignment of *HypBol_v2* and *HypBol_v1*. *Melitaea cinxia* present 31 Merian units, which represents the ancestral state of Ditrysia (Wright et al. 2023). *M. cinxia* belongs to the *Nymphalinae* which is the same subfamily as *Hypolimnas*, and the two taxa diverged about 30 MYa (Espeland et al. 2018). Merian units in *HypBol_v1* and *H. missipus* were assigned by choosing the chromosome sharing the highest BUSCO hits with a *M. cinxia* chromosome.

### Searches of synteny blocks

Synteny of the two assemblies was assessed using two methods. First, the two final genome assemblies were aligned using D-GENIES (Cabanettes & Klopp 2018), which produced a paf file that was used to detect candidate inversions between the two genome assemblies (Supplementary Figure 1). Second, synteny between the two assemblies was evaluated using Satsuma2 Synteny (Grabherr et al. 2010) and a circos plot generated using the circlize package v 0.4.14 (Gu et al. 2014) in R v 4.1.2 (Figure 2).

To examine the origin and evolution of the W chromosome in Lepidoptera, first, synteny-based homology between the *H. misippus* W chromosome, the Z chromosome and the autosomes was evaluated using Satsuma2. Satsuma2 is an aligner of whole genome assemblies intended to find homology based on sequence similarity. Satsuma2 first maps all genomic windows of the query genome to the target genome with a percentage of identity higher than 45%, and then filters those hits based on large scale synteny, that is, it keeps only matches that are concordant with each other. Thus, Satsuma2 is not only aligning genomic windows, but also evaluating synteny between those blocks. Finally, Satsuma2 focusses on regions with a high number of hits, to exhaustively evaluate the region around those hits, a strategy analogous to the battleship game. To evaluate synteny between the W and the remaining 31 chromosomes in *H. misippus,* the W chromosome was used as a query and the 31 assembled chromosomes as targets for Satstuma2. Resulting mapped regions were then filtered to keep only non-overlapping regions using the package GenomicRanges in R v4.1.2 (Lawrence et al. 2013). Satsuma2 does not require input genome assemblies to be masked. Because of its algorithm and filtering steps, repeat regions mapping to multiple places in the genome have decreased score and may be filtered out. However, to evaluate only the synteny of non-repeat regions, the RepeatMasker output, which details the coordinates of repeats in the genome, was used to filter out repeat regions of the W chromosome. Finally, the effect of sequence identity was evaluated by performing the analysis with no identity filter and applying a threshold of 70% similarity.

The synteny between the W and the rest of the genome was also evaluated in 6 other Lepidoptera species: *Boloria selene* (GCA_905231865.2), *Crocallis elinguaria* (GCA_907269065.1), *Hemaris fuciformis* (GCA_907164795.1), *Papilio machaon* (GCA_912999745.1), *Watsonalla binaria* (GCA_929442735.1), *Zygaena filipendulae* (GCA_907165275.2) and *Dryas iulia* (GCA_019049465.1). In all cases, the W chromosome was used as the query, results were converted to non-overlapping regions and percentage of the W covered by matches calculated. No filter based on repeats was applied, as no repeat library was available for these species.

After that, Satsuma2 was used to compare the synteny of the *H. misippus* W (used as query) and Z chromosomes to 10 Lepidoptera species, the 6 from the previous analysis and also *Hemaris fuciformis* (GCA_907164795.1), *Marasmarcha lunaedactyla* (GCA_923062675.1), *Vanessa cardui* (GCF_905220365.1), and *Mythimna farrago* (GCA_910589285.2). Chromosomes of all the species analysed were re-named by their homology to *M. cinxia* as above. Again, results were converted to non-overlapping regions, repeat regions of the W chromosome filtered and percentage of the W chromosome covered by matches calculated. Using only the *H. misippus* W chromosome as query ensures that secondary and tertiary matches are also reported. A more specific analysis was produced by using the *H. misippus* assembly as query to search for synteny in the 10 Lepidoptera target species. Using whole assemblies as input limits the results to only primary matches, pairing all homologous autosomes.

### BLAST of *H. misippus* W chromosome protein-coding genes

To further explore the degree of homology between the *H. misippus* W chromosome and the autosomes and Z chromosome, protein sequence homology was assessed using BLASTp v2.4.0+ (BLAST 2013; Kent 2002). First a protein BLAST database was built using the protein sequences contained in the *H. misippus* autosomes and the Z chromosome in the HypMisi_v2 annotation. Then, protein sequences found in the W chromosome were extracted and blasted to the database setting the minimum e-value to 1e-10 and the maximum number of target sequences to be reported to 5,000. The best match for each protein sequence was selected based on e-value score, and if two matched had the same value percentage of identity was used. Finally, genome coordinates were extracted from the HypMisi_v2 annotation.

## Acknowledgements

This work was supported by the Natural Environment Research Council [grant number NE/L002507/1 to A.O; NE/N010434/1 to GH]; the Cambridge Trust [European Research Scholarship to A.O] and St. John’s College [Benefactors’ Scholarship to A.O]. SHM was supported by a Royal Society University Research Fellowship [grant number URF\R1\180682].

## Data availability

Genome assemblies and annotations for *Hypolimnas misippus* and *Hypolimnas bolina* are available NCBI BioProject accession PRJEB64523. HiFi PacBio and Illumina raw reads of the *H. misippus* trio binning samples are available NCBI BioProject accession PRJEB64523. The data underlying this article are available in 10.5281/zenodo.8172459. Code used for analyses can be found in https://github.com/annaorteu/Hypolimnas_genome_Wchr_evolution.

## References

Bachtrog D et al. 2011. Are all sex chromosomes created equal? Trends in Genetics. 27:350–357. doi: 10.1016/j.tig.2011.05.005.

Bachtrog D et al. 2014. Sex Determination: Why So Many Ways of Doing It? PLOS Biology. 12:e1001899. doi: 10.1371/journal.pbio.1001899.

Berner D, Ruffener S, Blattner LA. 2023. Chromosome-Level Assemblies of the Pieris mannii Butterfly Genome Suggest Z-Origin and Rapid Evolution of the W Chromosome. Genome Biology and Evolution. 15:evad111. doi: 10.1093/gbe/evad111.

Beukeboom LW, Perrin N. 2014. The Evolution of Sex Determination. Oxford University Press doi: 10.1093/acprof:oso/9780199657148.001.0001.

Bishop G, Ebdon S, Lohse K, Vila R. 2021. The genome sequence of the small tortoiseshell butterfly, Aglais urticae (Linnaeus, 1758). doi: 10.12688/wellcomeopenres.17197.1.

BLAST. 2013. BLAST Basic Local Alignment Search Tool. doi: 10.1006/jmbi.1990.9999.

Brien MN et al. 2022. Colour polymorphism associated with a gene duplication in male wood tiger moths. 2022.04.29.490025. doi: 10.1101/2022.04.29.490025.

Brůna T, Hoff KJ, Lomsadze A, Stanke M, Borodovsky M. 2021. BRAKER2: automatic eukaryotic genome annotation with GeneMark-EP+ and AUGUSTUS supported by a protein database. NAR Genomics and Bioinformatics. 3:lqaa108. doi: 10.1093/nargab/lqaa108.

Cabanettes F, Klopp C. 2018. D-GENIES: dot plot large genomes in an interactive, efficient and simple way. PeerJ. 6:e4958. doi: 10.7717/peerj.4958.

Challis R, Richards E, Rajan J, Cochrane G, Blaxter M. 2020. BlobToolKit – Interactive Quality Assessment of Genome Assemblies. G3 Genes|Genomes|Genetics. 10:1361–1374. doi: 10.1534/g3.119.400908.

Charlat S et al. 2009. The joint evolutionary histories of Wolbachia and mitochondria in Hypolimnas bolina. BMC Evolutionary Biology. 9:64. doi: 10.1186/1471-2148-9-64.

Cheng H, Concepcion GT, Feng X, Zhang H, Li H. 2020. Haplotype-resolved de novo assembly with phased assembly graphs. arXiv:2008.01237 [q-bio]. http://arxiv.org/abs/2008.01237 (Accessed August 27, 2020).

Dai W, Mank JE, Ban L. 2022. Repeated origin of the W chromosome from the Z chromosome in Lepidoptera. 2022.10.21.512844. doi: 10.1101/2022.10.21.512844.

Dalíková M et al. 2017. New Insights into the Evolution of the W Chromosome in Lepidoptera. Journal of Heredity. 108:709–719. doi: 10.1093/jhered/esx063.

Dobin A et al. 2013. STAR: ultrafast universal RNA-seq aligner. Bioinformatics. 29:15–21. doi: 10.1093/bioinformatics/bts635.

Dudchenko O et al. 2017. De novo assembly of the Aedes aegypti genome using Hi-C yields chromosome-length scaffolds. Science. 356:92–95. doi: 10.1126/science.aal3327.

Dudchenko O et al. 2018. The Juicebox Assembly Tools module facilitates de novo assembly of mammalian genomes with chromosome-length scaffolds for under $1000. bioRxiv. 254797. doi: 10.1101/254797.

Durand NC, Robinson JT, et al. 2016. Juicebox Provides a Visualization System for Hi-C Contact Maps with Unlimited Zoom. Cell Systems. 3:99–101. doi: 10.1016/j.cels.2015.07.012.

Durand NC, Shamim MS, et al. 2016. Juicer Provides a One-Click System for Analyzing Loop-Resolution Hi-C Experiments. cels. 3:95–98. doi: 10.1016/j.cels.2016.07.002.

Dyson EA, Kamath MK, Hurst GDD. 2002. Wolbachia infection associated with all-female broods in Hypolimnas bolina (Lepidoptera: Nymphalidae): evidence for horizontal transmission of a butterfly male killer. Heredity. 88:166–171. doi: 10.1038/sj.hdy.6800021.

Espeland M et al. 2018. A Comprehensive and Dated Phylogenomic Analysis of Butterflies. Current Biology. 28:770–778.e5. doi: 10.1016/j.cub.2018.01.061.

Fraïsse C, Picard MAL, Vicoso B. 2017. The deep conservation of the Lepidoptera Z chromosome suggests a non-canonical origin of the W. Nat Commun. 8:1486. doi: 10.1038/s41467-017-01663-5.

Grabherr MG et al. 2010. Genome-wide synteny through highly sensitive sequence alignment: Satsuma. Bioinformatics. 26:1145–1151. doi: 10.1093/bioinformatics/btq102.

Gu Z, Gu L, Eils R, Schlesner M, Brors B. 2014. circlize implements and enhances circular visualization in R. Bioinformatics. 30:2811–2812. doi: 10.1093/bioinformatics/btu393.

Guan D. 2022. Purge_Dups. https://github.com/dfguan/purge_dups (Accessed September 29, 2022).

Hejníčková M et al. 2019. Absence of W Chromosome in Psychidae Moths and Implications for the Theory of Sex Chromosome Evolution in Lepidoptera. Genes. 10:1016. doi: 10.3390/genes10121016.

Hickey G, Paten B, Earl D, Zerbino D, Haussler D. 2013. HAL: a hierarchical format for storing and analyzing multiple genome alignments. Bioinformatics. 29:1341–1342. doi: 10.1093/bioinformatics/btt128.

Hornett EA et al. 2006. Evolution of male-killer suppression in a natural population. PLoS Biology. 4:1643–1648. doi: 10.1371/journal.pbio.0040283.

Kawahara AY et al. 2019. Phylogenomics reveals the evolutionary timing and pattern of butterflies and moths. Proc Natl Acad Sci U S A. 116:22657–22663. doi: 10.1073/pnas.1907847116.

Keilwagen J, Hartung F, Grau J. 2019. GeMoMa: Homology-Based Gene Prediction Utilizing Intron Position Conservation and RNA-seq Data. In: Gene Prediction: Methods and Protocols. Kollmar, M, editor. Methods in Molecular Biology Springer: New York, NY pp. 161–177. doi: 10.1007/978-1-4939-9173-0_9.

Kent WJ. 2002. BLAT - The BLAST-like alignment tool. Genome Research. doi: 10.1101/gr.229202. Article published online before March 2002.

Kolmogorov M et al. 2018. Chromosome assembly of large and complex genomes using multiple references. Genome Res. doi: 10.1101/gr.236273.118.

Kolmogorov M, Raney B, Paten B, Pham S. 2014. Ragout—a reference-assisted assembly tool for bacterial genomes. Bioinformatics. 30:i302–i309. doi: 10.1093/bioinformatics/btu280.

Krueger F. 2015. Trim galore. A wrapper tool around Cutadapt and FastQC to consistently apply quality and adapter trimming to FastQ files.

Lawrence M et al. 2013. Software for Computing and Annotating Genomic Ranges. PLOS Computational Biology. 9:e1003118. doi: 10.1371/journal.pcbi.1003118.

Lennuk L. 2023. Estonian Museum of Natural History Department of Zoology. doi: 10.15468/98CXTC.

Lewis JJ et al. 2021. The Dryas iulia Genome Supports Multiple Gains of a W Chromosome from a B Chromosome in Butterflies. Genome Biology and Evolution. 13. doi: 10.1093/gbe/evab128.

Li H. 2013. Aligning sequence reads, clone sequences and assembly contigs with BWA-MEM. arXiv:1303.3997 [q-bio]. http://arxiv.org/abs/1303.3997 (Accessed August 27, 2020).

Li H. 2022. lh3/yak. https://github.com/lh3/yak (Accessed September 25, 2022).

Li H et al. 2009. The Sequence Alignment/Map format and SAMtools. Bioinformatics. 25:2078–2079. doi: 10.1093/bioinformatics/btp352.

Lukhtanov VA. 2000. Sex chromatin and sex chromosome systems in nonditrysian Lepidoptera (Insecta). Journal of Zoological Systematics and Evolutionary Research. 38:73–79. doi: 10.1046/j.1439-0469.2000.382130.x.

Marsh NA, Clarke CA, Rothschild M, Kellett DN. 1977. Hypolimnas bolina (L.), a mimic of danaid butterflies, and its model Euploea core (Cram.) store cardioactive substances. Nature. 268:726–728. doi: 10.1038/268726a0.

MCZ Harvard University. 2023. Museum of Comparative Zoology, Harvard University. doi: 10.15468/P5RUPV.

Moore R. 2023. Auckland Museum Entomology Collection. doi: 10.15468/CADE8J.

Paten B et al. 2011. Cactus: Algorithms for genome multiple sequence alignment. Genome Res. 21:1512–1528. doi: 10.1101/gr.123356.111.

Picelli S et al. 2014. Tn5 transposase and tagmentation procedures for massively scaled sequencing projects. Genome Res. 24:2033–2040. doi: 10.1101/gr.177881.114.

Quick J. 2018. Ultra-long read sequencing protocol for RAD004. https://www.protocols.io/view/ultra-long-read-sequencing-protocol-for-rad004-mrxc57n (Accessed December 13, 2022).

Rastas P. 2020. Lep-Anchor: automated construction of linkage map anchored haploid genomes. Bioinformatics. 36:2359–2364. doi: 10.1093/bioinformatics/btz978.

Rastas P. 2017. Lep-MAP3: robust linkage mapping even for low-coverage whole genome sequencing data. Bioinformatics. 33:3726–3732. doi: 10.1093/bioinformatics/btx494.

Ruan J, Li H. 2020. Fast and accurate long-read assembly with wtdbg2. Nat Methods. 17:155–158. doi: 10.1038/s41592-019-0669-3.

Sahara K, Yoshido A, Traut W. 2012. Sex chromosome evolution in moths and butterflies. Chromosome Res. 20:83–94. doi: 10.1007/s10577-011-9262-z.

Sahoo RK et al. 2018. Evolution of Hypolimnas butterflies (Nymphalidae): Out-of-Africa origin and Wolbachia-mediated introgression. Molecular Phylogenetics and Evolution. 123:50–58. doi: 10.1016/j.ympev.2018.02.001.

Seppey M, Manni M, Zdobnov EM. 2019. BUSCO: Assessing Genome Assembly and Annotation Completeness. In: Gene Prediction: Methods and Protocols. Kollmar, M, editor. Methods in Molecular Biology Springer: New York, NY pp. 227–245. doi: 10.1007/978-1-4939-9173-0_14.

Simão FA, Waterhouse RM, Ioannidis P, Kriventseva EV, Zdobnov EM. 2015. BUSCO: assessing genome assembly and annotation completeness with single-copy orthologs. Bioinformatics. 31:3210–3212. doi: 10.1093/bioinformatics/btv351.

Singh KS et al. 2022. Genome assembly of Danaus chrysippus and comparison with the Monarch Danaus plexippus. G3 Genes|Genomes|Genetics. 12:jkab449. doi: 10.1093/g3journal/jkab449.

Smit A, Hubley R, Green P. 2015. RepeatMasker Open-4.0. 2013–2015.

Smith DAS. 1976. Phenotypic diversity, mimicry and natural selection in the African butterfly Hypolimnas misippus L. (Lepidoptera: Nymphalidae). Biological Journal of the Linnean Society. 8:183–204. doi: 10.1111/j.1095-8312.1976.tb00245.x.

Storer J, Hubley R, Rosen J, Wheeler TJ, Smit AF. 2021. The Dfam community resource of transposable element families, sequence models, and genome annotations. Mobile DNA. 12:2. doi: 10.1186/s13100-020-00230-y.

Turner JRG, Sheppard PM. 1975. Absence of crossing-over in female butterflies (Heliconius). Heredity. 34:265–269. doi: 10.1038/hdy.1975.29.

Tyson J. 2020. Bead-free long fragment LSK109 library preparation. https://www.protocols.io/view/bead-free-long-fragment-lsk109-library-preparation-7eshjee (Accessed December 13, 2022).

Vane-Wright RI, Ackery PR, Smiles RL. 1977. The polymorphism, mimicry, and host plant relationships of Hypolimnas butterflies. Biological Journal of the Linnean Society. 9:285–297. doi: 10.1111/j.1095-8312.1977.tb00271.x.

Vítková M, Fuková I, Kubíčková S, Marec F. 2007. Molecular divergence of the W chromosomes in pyralid moths (Lepidoptera). Chromosome Res. 15:917–930. doi: 10.1007/s10577-007-1173-7.

Walker BJ et al. 2014. Pilon: An Integrated Tool for Comprehensive Microbial Variant Detection and Genome Assembly Improvement. PLOS ONE. 9:e112963. doi: 10.1371/journal.pone.0112963.

Wick R. 2022. Porechop. https://github.com/rrwick/Porechop (Accessed September 28, 2022).

Wright AE, Dean R, Zimmer F, Mank JE. 2016. How to make a sex chromosome. Nat Commun. 7:12087. doi: 10.1038/ncomms12087.

Wright CJ, Stevens L, Mackintosh A, Lawniczak M, Blaxter M. 2023. Chromosome evolution in Lepidoptera. 2023.05.12.540473. doi: 10.1101/2023.05.12.540473.

Wu Y-W, Simmons BA, Singer SW. 2016. MaxBin 2.0: an automated binning algorithm to recover genomes from multiple metagenomic datasets. Bioinformatics. 32:605–607. doi: 10.1093/bioinformatics/btv638.

Wulf A. 2016. The Woman Who Made Science Beautiful. The Atlantic. https://www.theatlantic.com/science/archive/2016/01/the-woman-who-made-science-beautiful/424620/ (Accessed July 7, 2022).

Yang J et al. 2020. Chromosome-level reference genome assembly and gene editing of the dead-leaf butterfly Kallima inachus. Molecular Ecology Resources. 20:1080–1092. doi: 10.1111/1755-0998.13185.

Yen EC et al. 2020. A haplotype-resolved, de novo genome assembly for the wood tiger moth (Arctia plantaginis) through trio binning. Gigascience. 9. doi: 10.1093/gigascience/giaa088.

Yoshida K et al. 2011. B Chromosomes Have a Functional Effect on Female Sex Determination in Lake Victoria Cichlid Fishes. PLOS Genetics. 7:e1002203. doi: 10.1371/journal.pgen.1002203.

Yoshido A, Šíchová J, Kubíčková S, Marec F, Sahara K. 2013. Rapid turnover of the W chromosome in geographical populations of wild silkmoths, Samia cynthia ssp. Chromosome Res. 21:149–164. doi: 10.1007/s10577-013-9344-1.

