## Supplementary data for "The *Hypolimnas misippus* genome supports a common origin of the W chromosome in Lepidoptera"

**Supplementary Material**

| Feature | *HypBol_v1* | *HypMisi_v2* | *Vanessa cardui* |
|---|---|---|---|
| Total length: Gene | 160709320 | 167739620 | 240814091 |
| Total length: mRNA | 206176689 | 222209628 | 522618316 |
| Total length: CDS | 29265705 | 32999245 | 38521385 |
| Total length: Exon | 35299426 | 39816515 | 56622152 |
| Total length: Intron | 170992076 | 182514332 | 465996164 |
| Mean number per mRNA: Exons | 6.3 | 6.4 | 9.5 |
| Mean number per mRNA: Introns | 5.3 | 5.4 | 8.5 |
| Mean length: Gene | 8149 | 8265 | 18211 |
| Mean length: mRNA | 9464 | 9890 | 26346 |
| Mean length: Exon | 258 | 277 | 299 |
| Mean length: Intron | 1489 | 1505 | 2754 |
| Total number: Genes | 19721 | 20293 | 13223 |
| Total number: mRNAs | 21784 | 22468 | 19836 |
| Total number: Exons | 136597 | 143687 | 189042 |
| Total number: Introns | 114813 | 121219 | 169206 |
| Total number: Single exon genes | 3868 | 4891 | 1387 |
| % of genome covered by: Genes | 36.1 | 38.3 | 56.7 |
| % of genome covered by: mRNAs | 36.1 | 38.2 | 54.4 |
| % of genome covered by: Exons | 5.6 | 6.3 | 7.2 |
| % of genome covered by: Introns | 30.5 | 32 | 47.1 |

**Supplementary Table 1.** Annotation statistics of *Hypolimnas misippus (HypMisi_v2)* and *H. bolina (HypBol_v1)* compared to *Vanessa cardui*.

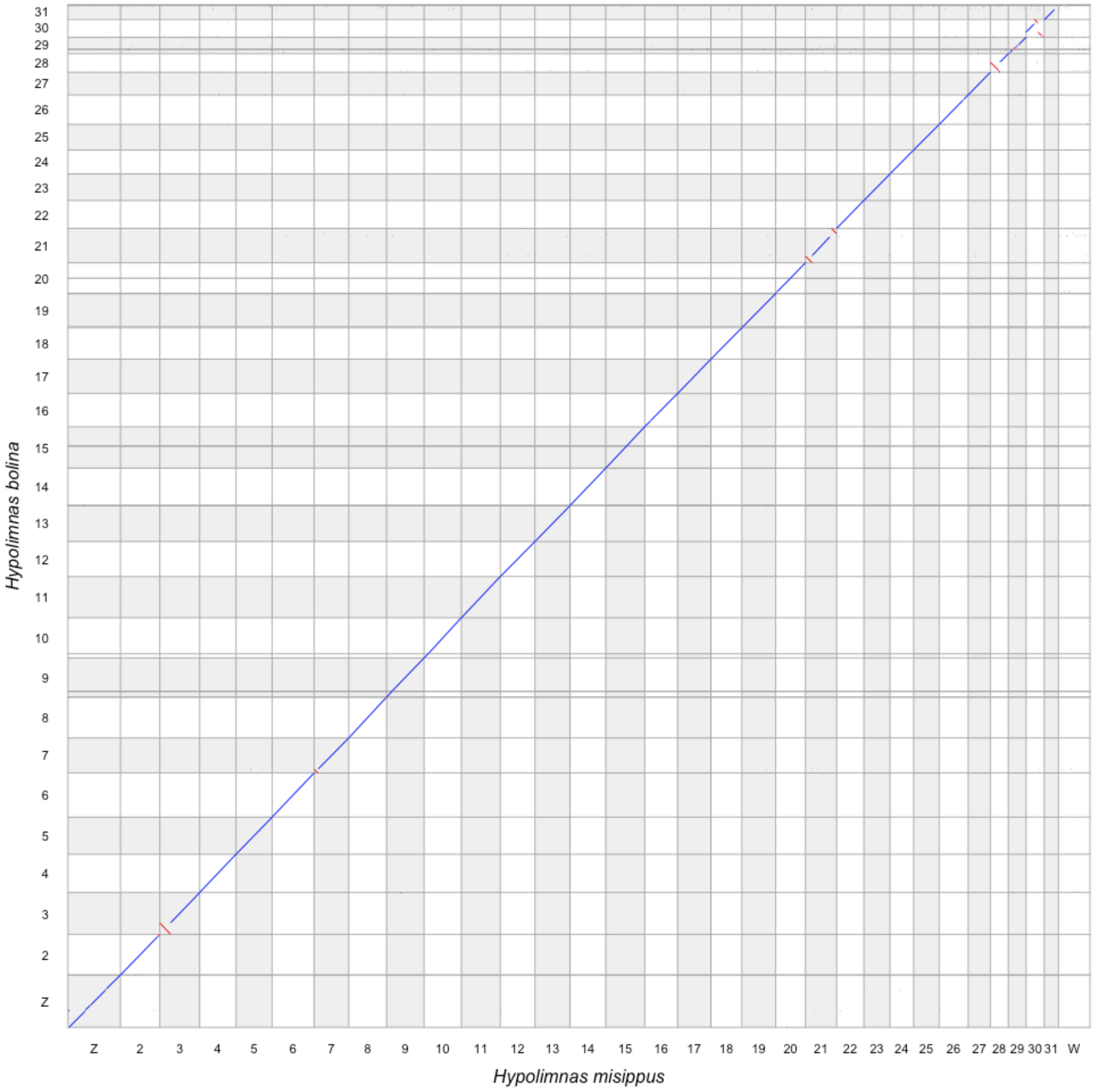


**Supplementary Figure 1.** The D-Genies whole assembly alignment of *Hypolimnas misippus* and *H. bolina* reveals 12 inversions between the two genomes (shown in red).

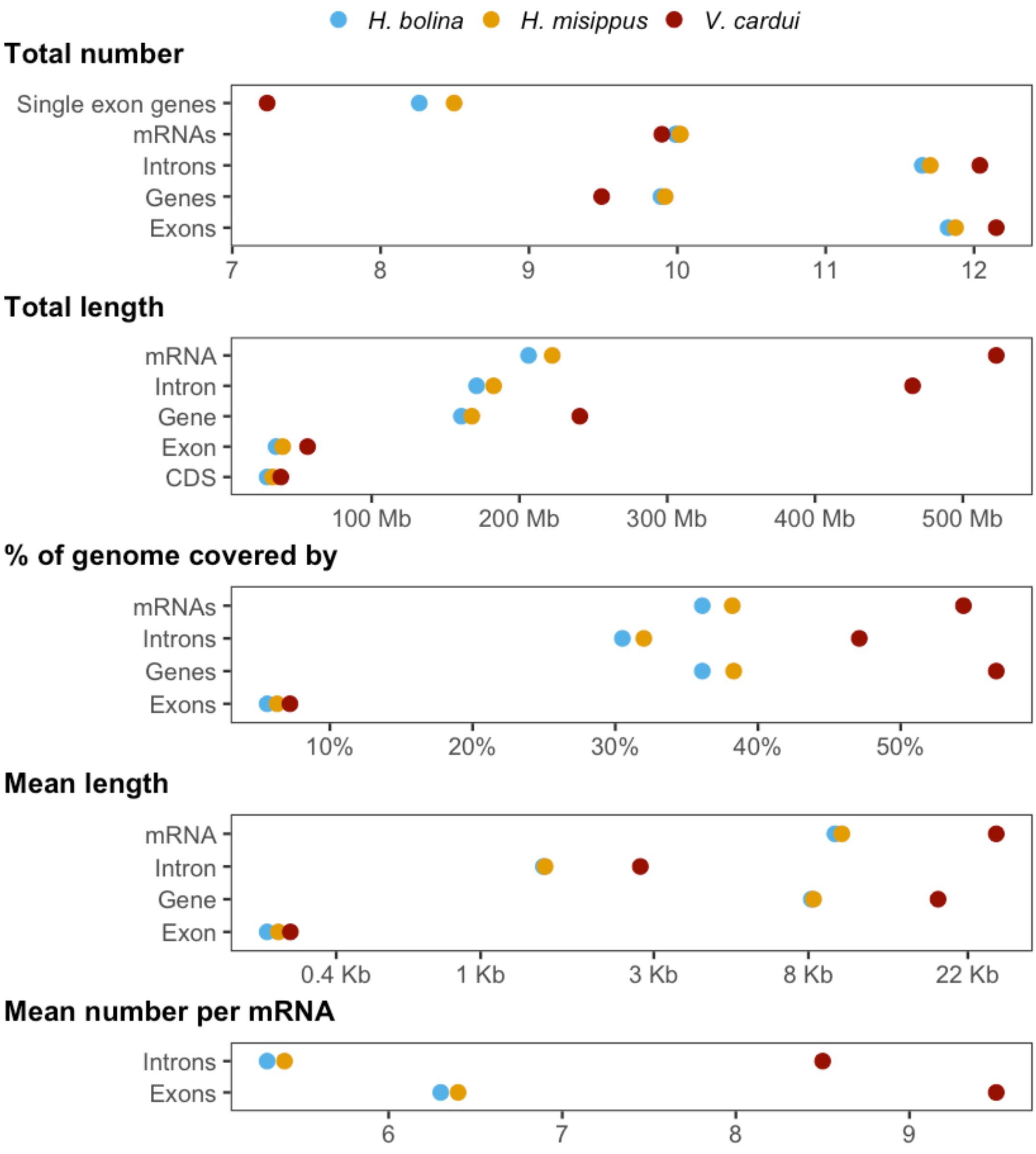


**Supplementary Figure 2.** Annotation comparison of *Hypolimnas misippus, H. bolina* and *Vanessa cardui*. For the statistics, pre-mRNA, that is mRNA including exons and introns, have been used. Numbers are found in Supplementary Table 1.

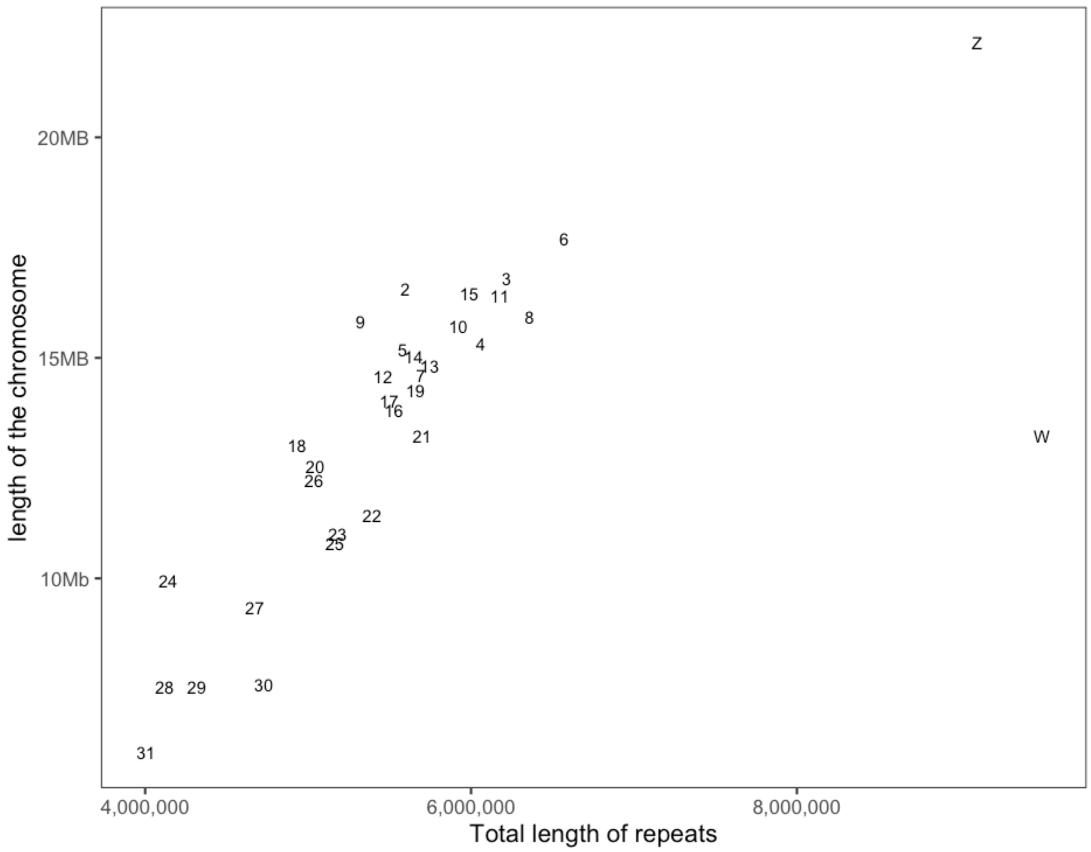


**Supplementary Figure 3.** Repeat content of the *HypMisi_v2* by chromosome. Larger chromosomes tend to have more repeats. The W chromosome deviates from the correlation and shows a higher repeat content for its length (see also Supplementary Figure 4).

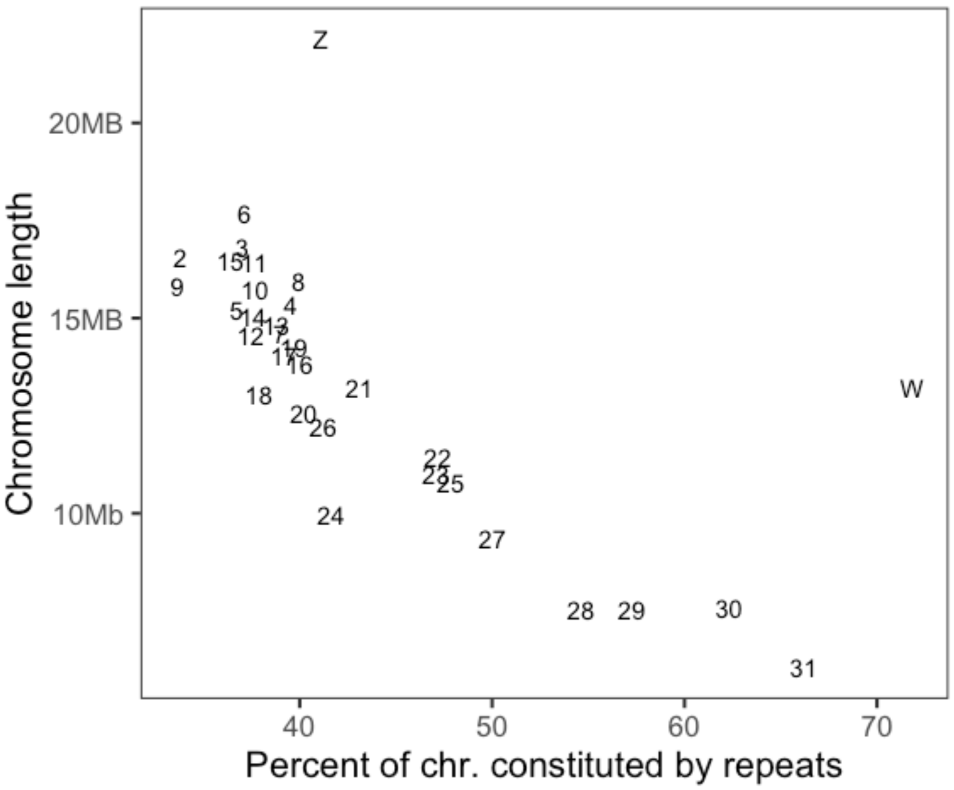


**Supplementary Figure 4.** Percentage of each chromosome of the *HypMisi_v2* assembly constituted by repeats. The W chromosome has the highest percentage of repeats.

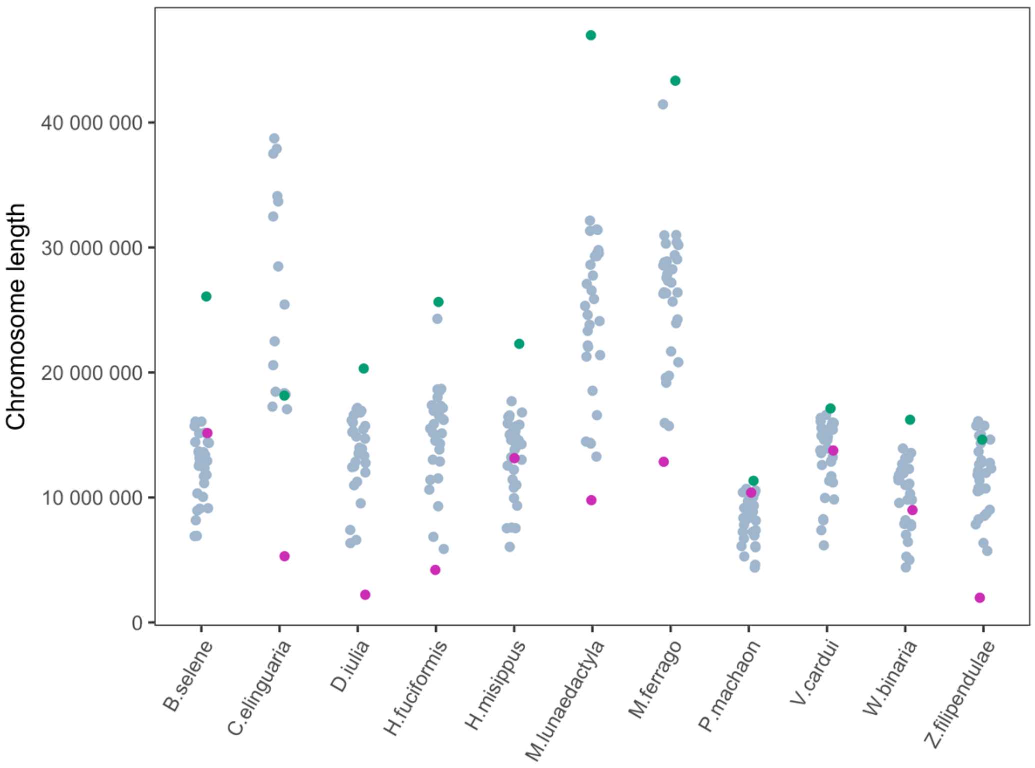


**Supplementary Figure 5.** Chromosomes sizes of the 11 Lepidoptera species used for the comparisons. Autosomes are shown in grey, W chromosomes in pink and Z chromosomes in green.

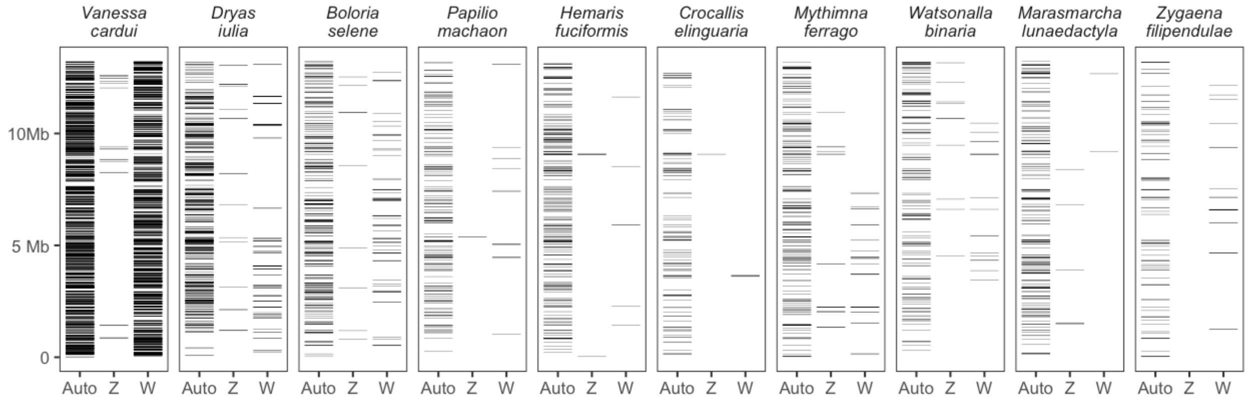


**Supplementary Figure 6.** The positions of synteny blocks between the *H. misippus* W chromosome and the chromosomes of other species do not reveal a conserved section of the W chromosome across species. Left-most column shows all synteny blocks with autosomes plotted.

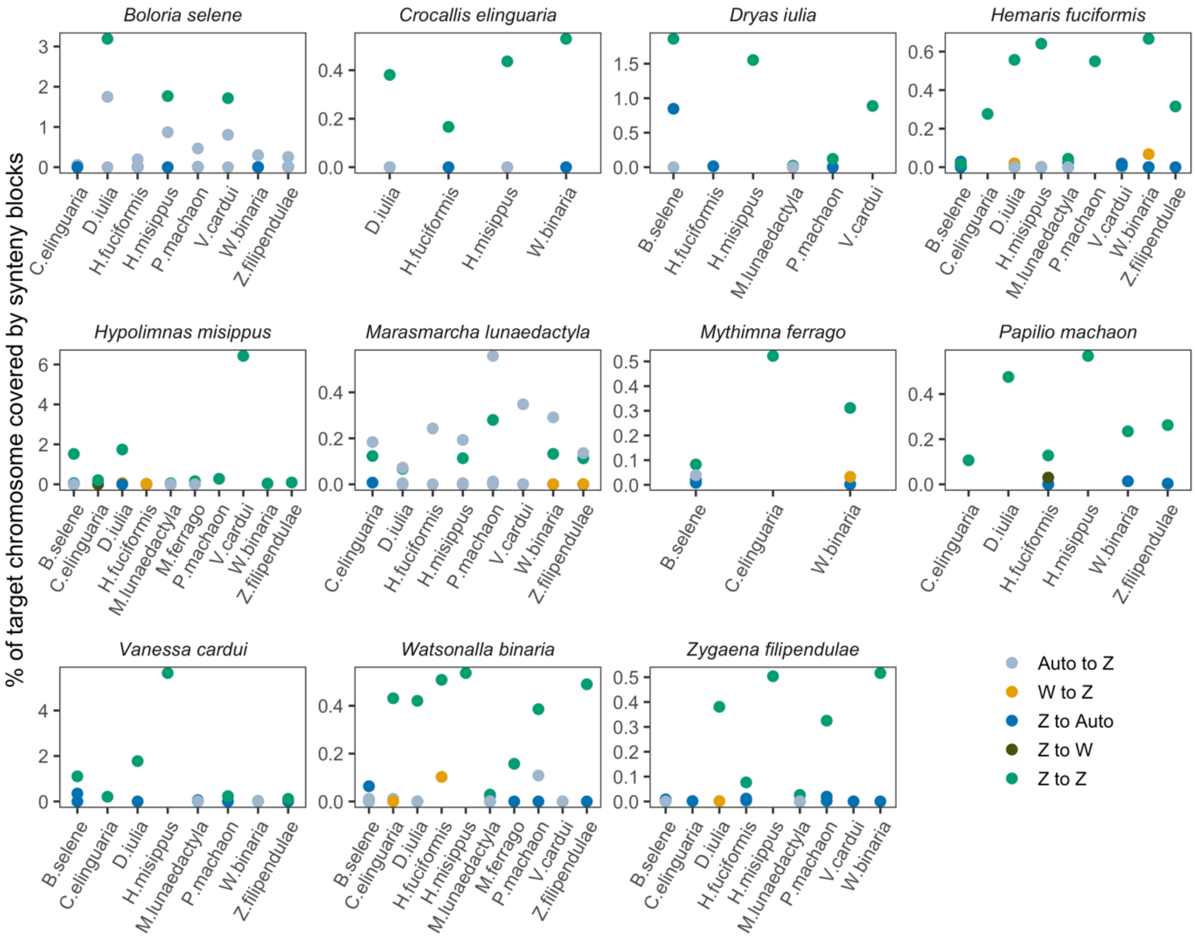


**Supplementary Figure 7.** Synteny analysis using Satsuma2 between 11 ditrysian species reveals deep conservation of the Z chromosome across species. Auto refers to any autosomes.

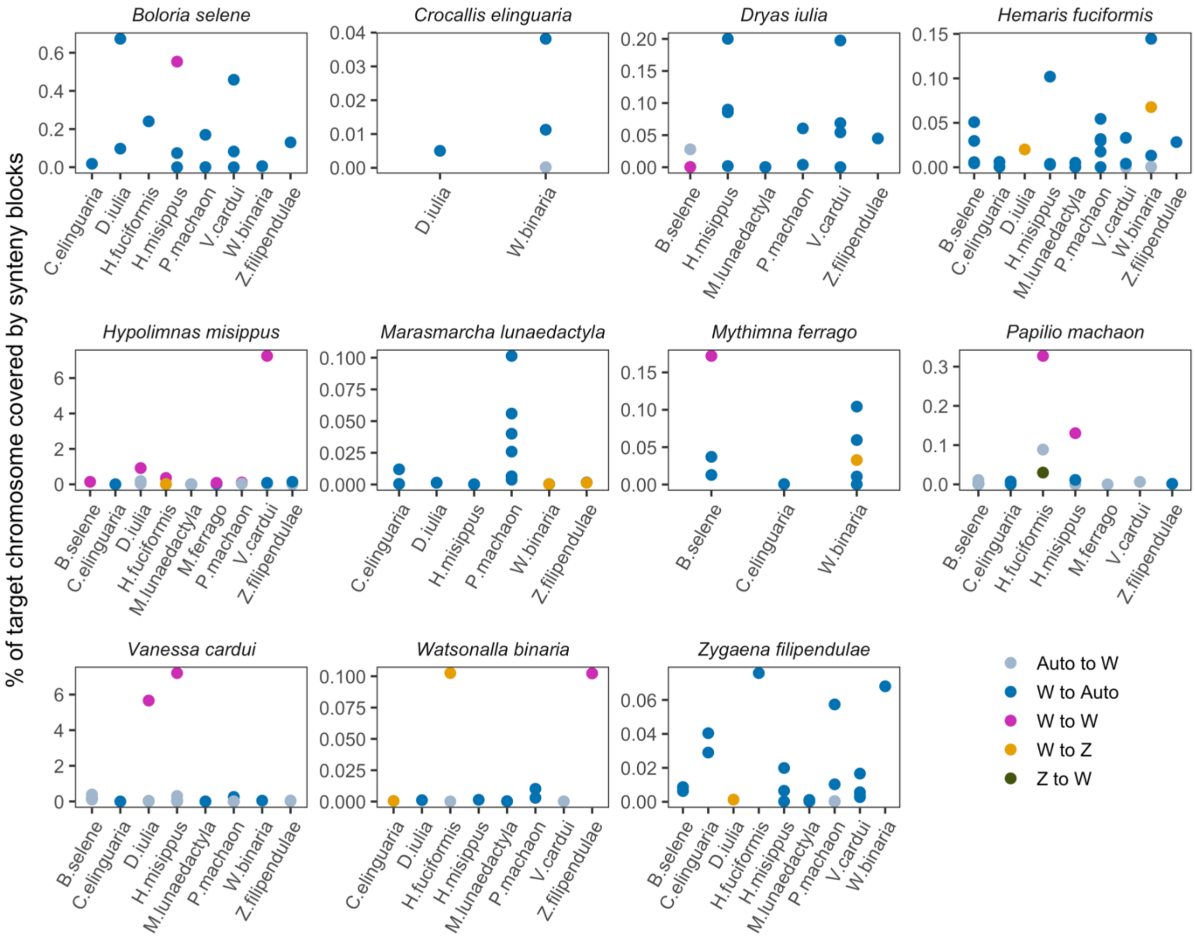


**Supplementary Figure 7.** Synteny analysis using Satsuma2 between 11 ditrysian species reveals some conservation of the W chromosome across species. Auto refers to any autosomes.

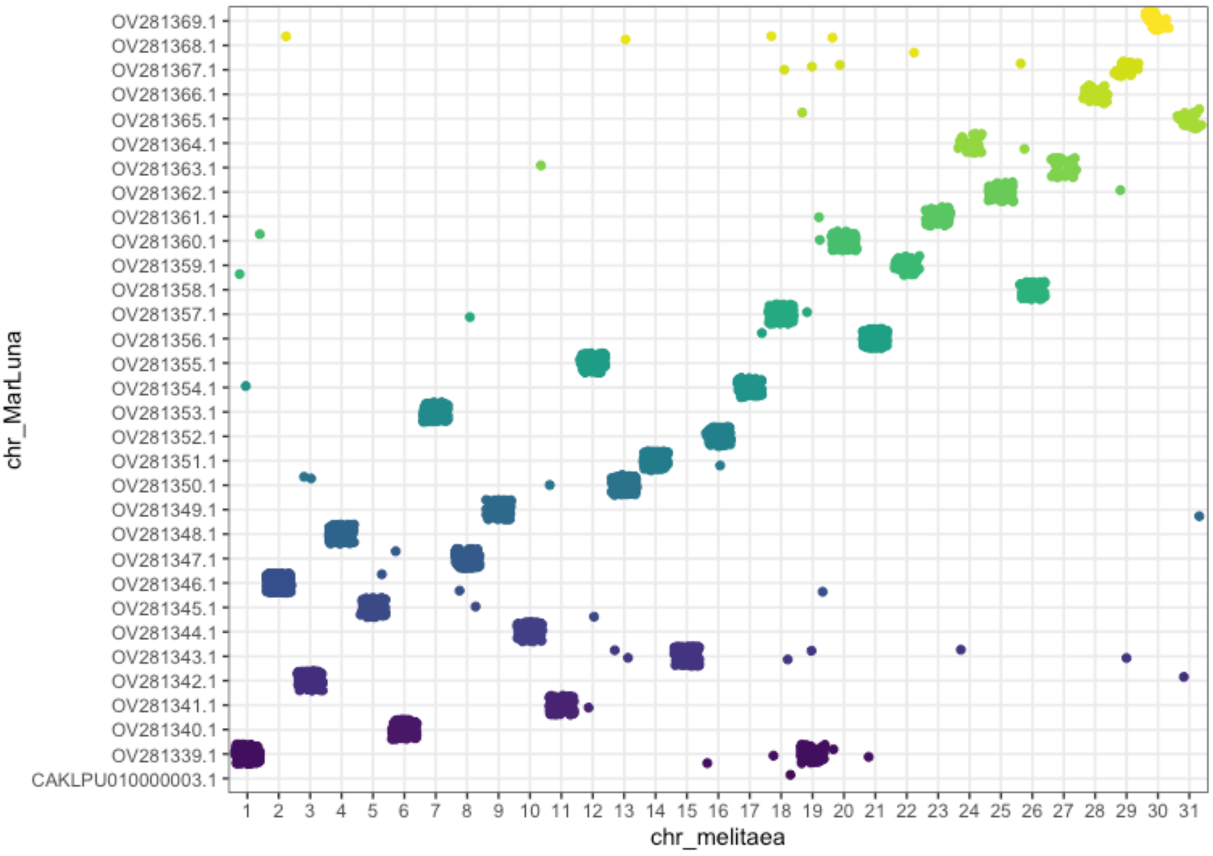


**Supplementary Figure 9.** Homology of *M. lunaedactyla's* chromosomes with *Melitaea cinxia* reveals a neo-Z chromosome. Chromosome OV281339.1 of *M. lunaedactyla* shares BUSCOs with chromosome 1 (Z) and chromosome 19 of *M. cinxia*, suggesting a fusion of these two.

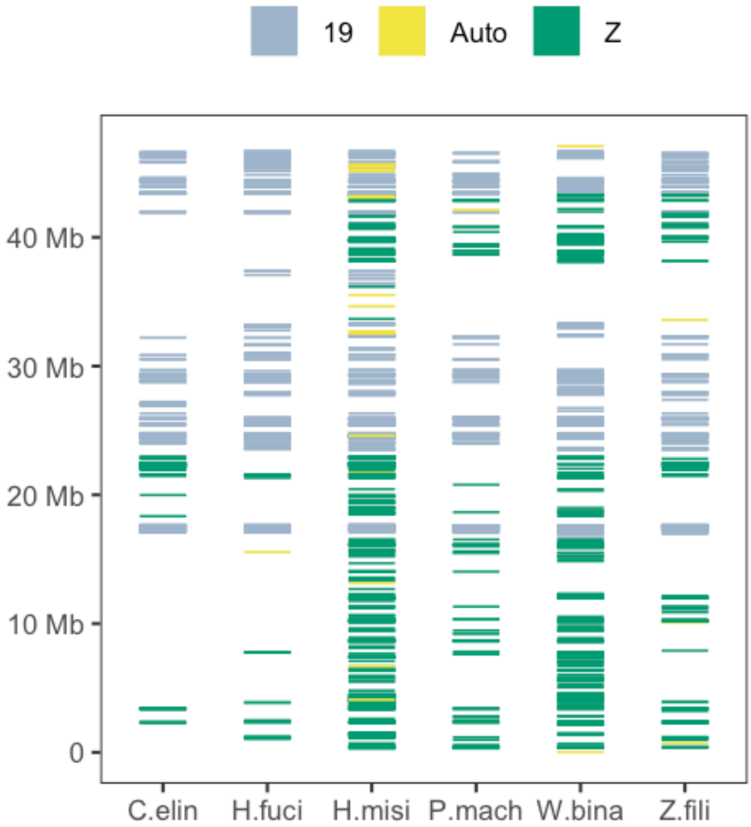

**Supplementary Figure 10.** Synteny analysis using Satsuma2 of the neo-Z chromosome of *M. lunaedactyla* compared to other ditrysian assemblies reveals the patterns of homology with the ancestral Z and chromosome 19.
